# Biosynthetic additive manufacturing through mycelium-bacteria consortia

**DOI:** 10.64898/2026.10.29.762868

**Authors:** Hang Yuan, Linux Joseph Jones Heller, Yi Xing, Lizhe Wang, Jiawei Huang, Khoi Nguyen, Yulu Cai, Bryce Waller, Cheng Chen, Grant Hubbard, Ruiqi Yong, Eleanor DelPlace, Miranda Belo, Suhani Manosh, Soham Inamdar, Xiwei Shan, Xiaodong Zang, Juhua Huang, Tian Li, Gregory Bonito, Kai Lan, Jinxing Li

## Abstract

Lignocellulosic biomass is the most abundant renewable raw material on Earth, yet its structural and carbon-storage potential remains unrealized: microbial decomposition returns stored carbon to the atmosphere before it can be harnessed as a durable material. Distinct microbial processes, such as biosynthesis and biomineralization, offer a natural counterpoint, converting organic carbon into mechanically robust, persistent materials such as polymers and minerals. Here, we establish a biosynthetic additive manufacturing strategy through mycelium–bacteria consortia, by coupling fungal self-assembly with bacterial biomineralization inside fabricated three-dimensional scaffolds derived from lignocellulosic biomass. *Trametes versicolor* forms a continuous living mycelial network that serves as both a structural template and a transport pathway, guiding the spatial colonization of *Sporosarcina pasteurii* and enabling volumetric calcium carbonate deposition throughout the scaffold interior, overcoming the surface-limited mineralization of conventional microbial systems. Integration with helicoidal Bouligand architectures translates microscale mineral reinforcement into macroscale performance, increasing compressive Young’s modulus by 7-14-fold and ultimate strength by 5-fold relative to unmineralized conventional mycelium-based living composites. The living mycelial network also enables autonomous self-healing and modular self-assembly through hyphal bridging without external adhesives or energy input. This work establishes a scalable route for converting and upcycling transient biomass carbon into durable living structural materials for sustainable construction.

## INTRODUCTION

The built environment generates 40% of annual global CO_2_ emissions, while building and infrastructure materials and construction (typically referred to as embodied carbon) are responsible for 13% annually, driven by reliance on energy-intensive materials (*1*, *2*). Lignocellulosic biomass, as the most abundant renewable raw material on Earth, offers a highly attractive alternative because of its high specific strength, low carbon footprint, and widespread availability (*3*, *4*). Large quantities of underutilized biomass, including fallen wood and agricultural residues, function as major carbon reservoirs in natural ecosystems, storing more than 73 billion tons of carbon globally (*5*, *6*). Yet this reservoir is fundamentally transient. Microbial decomposers continuously return stored carbon to the atmosphere, ensuring that most biomass participates in a short-term carbon cycle rather than serving as a durable structural sink (*7*).

The variety of microbial processes provides an alternative pathway. Unlike microbial decomposition, through microbial synthesis and biomineralization, carbon can be converted into mechanically strong and persistent materials such as cellulose networks, calcium carbonate, and other biominerals across multiple length scales, from nanocrystals to reef-scale stromatolites (*8–10*). Such microbial biosynthetic processes play significant roles in terraforming the planet and provide sinks for atmospheric carbon and other elements through biogeochemical cycling (*11*, *12*). In some wood-associated ecosystems, microbial activity reinforces rather than degrades the substrate by inducing localized mineral deposition that improves hardness and durability (*13*). These observations reveal a fundamental but underexploited principle: microbial metabolisms can be redirected from decomposition toward structural reinforcement, enabling biomass to function not only as a renewable feedstock but as a platform for material synthesis and long-term carbon sequestration.

Recent advances in engineered living materials have demonstrated the potential of biology-driven materials reuse and fabrication (*14*, *15*). Mycelium, the filamentous network of fungi, has emerged as an attractive platform because of its ability to self-assemble, biodegrade, and colonize substrates derived from agricultural and industrial waste (*16*, *17*). These properties enable the formation of lightweight porous composites with tunable mechanical and thermal characteristics, leading to applications in packaging, insulation, and construction (*18–21*). However, the inherently porous mycelial architecture still limits load-bearing performance, while existing systems lack a mechanism to coordinate structural reinforcement throughout the bulk material after fabrication.

Here, we introduce a biosynthetic additive manufacturing strategy that resolves these limitations by coupling three-dimensional (3D) printing, fungal self-assembly, and bacterial biomineralization within a unified fabrication framework. Lignocellulosic biomass, formulated as a printable bioink, serves simultaneously as a structural matrix, carbon reservoir, and microbial growth substrate. Within helicoidal Bouligand scaffolds, *Trametes versicolor* (*T. versicolor*) establishes a continuous, hierarchical hyphal network that functions as a 3D biological highway: guiding spatial colonization by the calcifying bacterium *Sporosarcina pasteurii* (*S. pasteurii*), supporting nutrient transport, and providing nucleation sites for *in situ* calcium carbonate deposition throughout the scaffold volume. By coupling architected 3D printing with controlled dual-microbial growth and mineralization, we transform waste biomass into an organic–inorganic living composite with volumetric mineral reinforcement, achieving a 7-14-fold increase in compressive Young’s modulus and a 5-fold increase in ultimate strength over unmineralized mycelium composites, while improving the specific strength of mycelium-based biocomposites as lightweight materials. Beyond static reinforcement, the living nature of the mycelial network enables autonomous self-healing of damaged interfaces and modular self-assembly of discrete printed units through hyphal bridging, without external adhesives or additional fabrication steps. This work establishes a new paradigm of biosynthetic additive manufacturing, in which material formation arises from the coupling of digital fabrication and biological growth, offering a scalable pathway toward carbon-negative, self-organizing structural materials for construction and beyond.

## RESULTS

### Dual-microbial biosynthetic additive manufacturing

One of the key challenges in engineered living materials is to ensure that biological growth strengthens rather than compromises structural construction (*22*, *23*). Unlike conventional bioprinting, where biology is treated as passive cargo, here biology serves as the construction mechanism. Therefore, we developed a dual-microbial fabrication strategy that integrates mycelium and biomineralizing bacteria within a 3D-printed scaffold, where each biological stage enables the next (**Fig. 1A**). Fabrication begins with a lignocellulosic biomass-derived bioink pre-colonized by *T. versicolor* mycelium and rheologically tuned for extrusion while preserving fungal viability. Immediately after printing, the scaffold serves only as a geometric template. During incubation, hyphae extend from printed filaments, bridge adjacent filaments and layers, and establish a continuous, living, 3D network. This mycelial network converts discrete printed lines into a coherent load-bearing scaffold and creates internal biological pathways that enable volumetric mineralization rather than surface-confined deposition (*24*). The scaffold is then co-cultured with *S. pasteurii*, a ureolytic bacterium capable of microbially induced calcium carbonate precipitation (MICP) (*25*, *26*). Instead of remaining confined to exposed surfaces, the pre-established hyphal network enables bacterial colonization throughout the scaffold interior (*25*). Urea hydrolysis generates local carbonate supersaturation (*27*), while the hyphal surface provides attachment and nucleation sites that localize mineral formation directly along the structural framework. The mycelium, therefore, serves simultaneously as a geometric template, transport network, and mineralization scaffold, transforming MICP from a surface-coating process into a volumetric reinforcement mechanism.

**Fig. 1.**
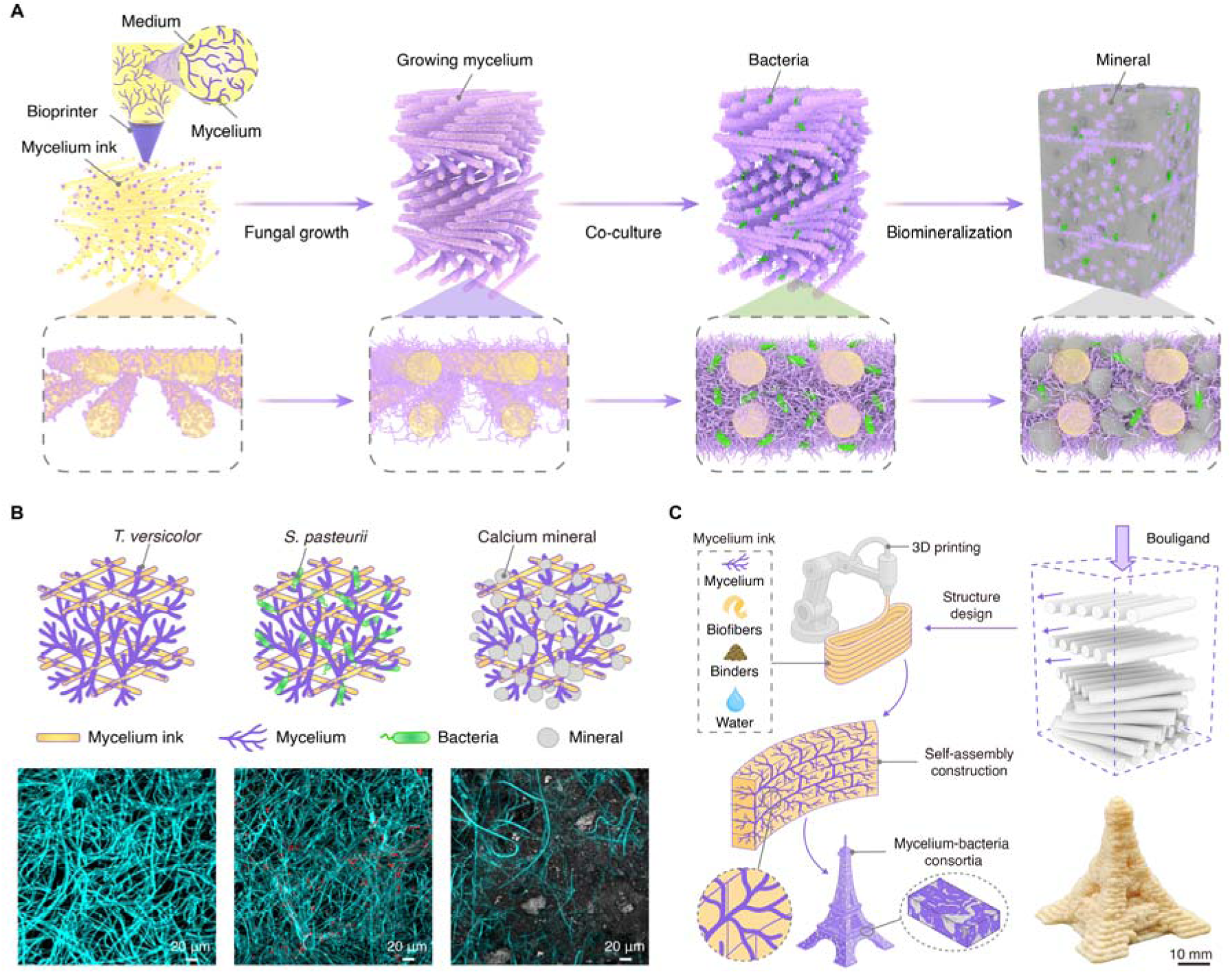
Dual-microbial biosynthetic additive manufacturing for volumetric structural reinforcement. (**A**) A stepwise schematic of the living composite biofabrication process. The process begins with the 3D printing of mycelium ink into a predefined scaffold. Subsequent fungal growth establishes a mycelial network. Co-culturing with biomineralizing bacteria (*S. pasteurii*) induces *in situ* calcium carbonate deposition within the network, forming a mineralized composite. (**B**) CLSM images of mycelial scaffolds at different stages. Left: Mycelial network formed by *T. versicolor* (shown in cyan). Middle: Mycelial network co-cultured with *S. pasteurii*, exhibiting fungal hyphae and membrane-compromised bacterial cells (shown in red). Right: Mineralized scaffold with fungal hyphae and calcium deposits (shown in gray). The images demonstrate the progressive colonization of the mycelial network by bacteria and the *in situ* formation of calcium minerals. (**C**) Potential of biosynthetic additive manufacturing for building construction. The ink composition (mycelium, biofibers, binders, and water) enables 3D printing of designed structures such as helicoidal stacking structures (Bouligand), followed by mycelial growth for self-assembly and initiation of biomineralization, leading to mycelium–bacteria consortia with enhanced structural performance. Bottom right: An Eiffel Tower model printed with MEA-based mycelium ink.

Confocal laser scanning microscopy (CLSM) confirmed this spatial organization at the microscale (**Fig. 1B**). In mycelium-only scaffolds, *T. versicolor* formed dense and continuous hyphal networks. In co-cultured scaffolds, *S. pasteurii* localized along hyphal intersections and branching points, and calcium-associated fluorescence signals appeared in regions closely associated with the hyphal network, consistent with *in situ* calcium carbonate deposition along biological pathways rather than as external bulk precipitates. To translate microscale mineral reinforcement into macroscale load-bearing performance, we introduced a Bouligand architecture with controlled interlayer rotation (**Fig. 1C**). This helicoidal geometry improves crack deflection and stress redistribution under compression while preserving the open interlayer pathways required for vertical hyphal bridging across layers (*25*). Architecture and biology are therefore co-designed: the geometry that best resists mechanical failure is also the geometry that best supports biological colonization and distributed mineralization. The printed construct thus evolves from a passive scaffold into an architected living composite, where final mechanical performance is determined not at the moment of printing, but through the biological activity that follows.

### Printable biomass-derived mycelium-inoculated living inks

A printable living scaffold needs to satisfy two competing requirements: reliable extrusion and sustained fungal growth after deposition (*28*). Inks optimized for printability often rely on synthetic polymers incompatible with long-term biological activity, whereas nutrient-rich media that support fungal growth typically lack the rheological stability needed for shape retention. To accommodate these competing demands, we developed three representative mycelium inks using sawdust-based biomass matrices, malt extract agar (MEA), and a commercially available Pluronic F127 (F127) (**Fig. 2A**) (*29*, *30*). These formulations were intentionally selected to represent three distinct roles in living biofabrication. MEA provides a nutrient-rich biological medium that supports rapid fungal growth, but its readily accessible nutrients also promote bacterial contamination during extended cultivation, limiting its long-term use as a structural material (*15*). F127 serves as a synthetic thermoresponsive hydrogel control with excellent extrusion behavior and immediate print fidelity; however, its temperature-dependent gelation makes it unsuitable for long-term cultivation and structural applications because the scaffold gradually loses stability during prolonged incubation (*31*). In contrast, sawdust functions simultaneously as a printable structural matrix, a carbon reservoir, and a metabolizable substrate for fungal growth (*32*, *33*). As a lignocellulosic biomass, it allows fungal colonization while maintaining dimensional stability, making it more suitable for living structural materials than temporary hydrogel carriers (*13*, *34*). Fungal species selection followed the same principle: the fungus must not only grow, but grow in a way that contributes to structural function (*35*). *T. versicolor* showed a good combination of rapid radial expansion, strong hyphal bridging, and stable colonization within printed biomass matrices. As a white-rot fungus with efficient lignocellulose degradation ability, *T. versicolor* rapidly penetrates biomass-derived scaffolds and forms dense filamentous networks across adjacent printed filaments and layers, which contributes to both scaffold continuity and subsequent mechanical reinforcement (*32*, *36*). Although *Ganoderma sessile* (*G. sessile*) also showed successful growth, *T. versicolor* demonstrated faster colonization and more reliable bridging performance (*36*, *37*), and was therefore selected as the primary fungal species for subsequent bioink development. All formulations showed good printability and enabled continuous filament extrusion (**Fig. 2B**), allowing fabrication of 3D structures that maintained geometry while supporting fungal growth after 7 days of incubation (**Fig. 2C**, fig. S1, and movies S1-2). This confirms that the printing process preserves fungal viability and allows the scaffold to transform from a printed object into a functional living network after fabrication rather than during extrusion. All inks exhibited shear-thinning behavior suitable for extrusion printing (**Fig. 2D**). For grid-like scaffolds, printed filaments must maintain sufficient storage modulus (*G’*) and yield stress (T_y_) to resist deformation caused by gravity and capillary forces (*15*, *38*). The sawdust-based mycelium ink showed the highest yield stress (565 Pa) and storage modulus (**Fig. 2E**), providing superior shape retention and deformation resistance. It also displayed the strongest structural recovery after repeated shear cycles (**Fig. 2F**), indicating better long-term stability during cultivation. In contrast, F127- and MEA-based inks showed weaker structural stability (fig. S2).

**Fig. 2.**
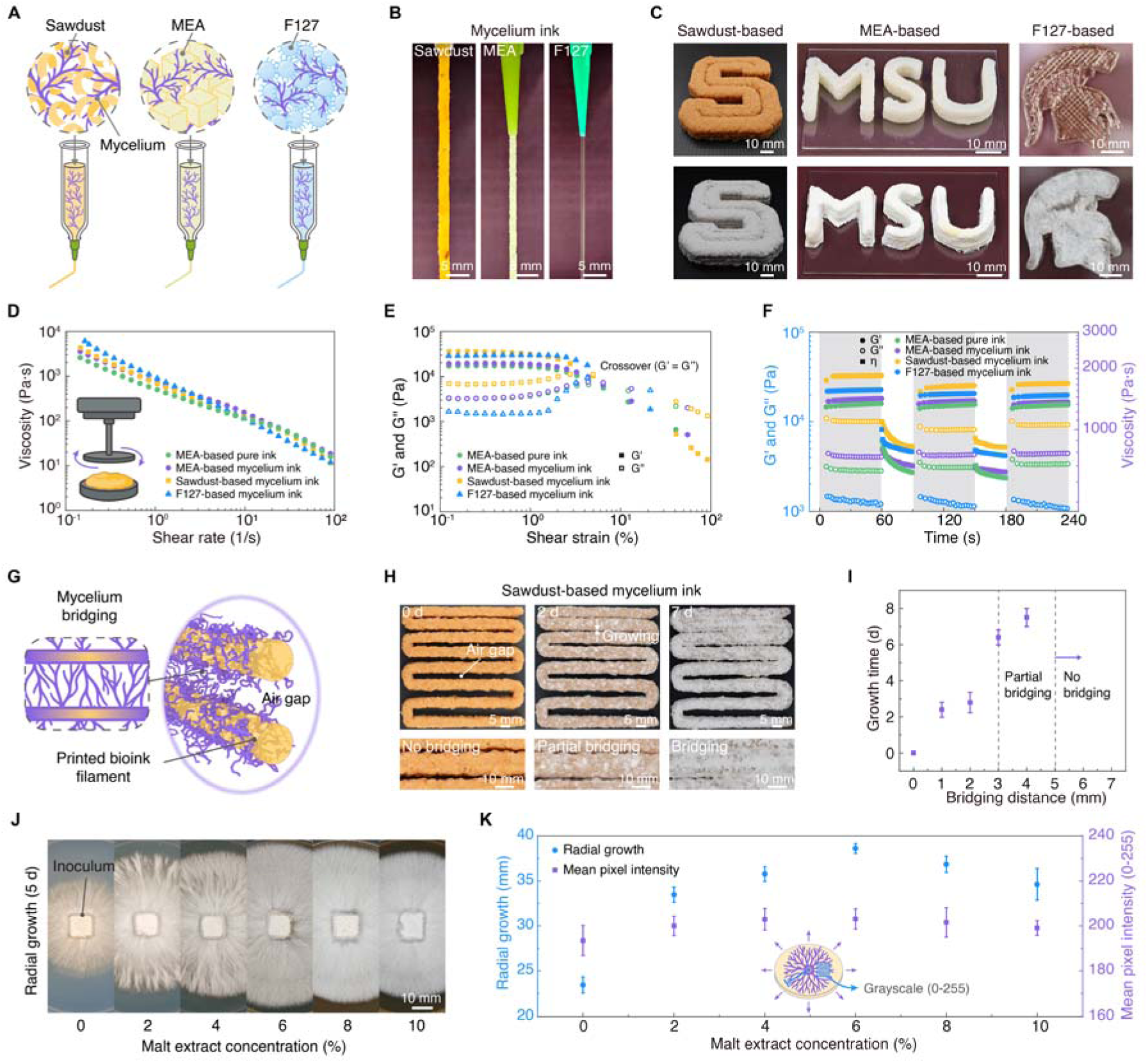
Rheological properties and printability of living mycelium inks prepared with different media and fungal growth behavior. (**A**) Schematic of mycelium inks formed by inoculating fungal mycelia into sawdust, MEA and F127 matrix. (**B**) Demonstration of the printability of sawdust-, MEA-, and F127-based mycelium inks. All formulations can be extruded through the printing nozzle into continuous filaments of centimeter-scale length. (**C**) Structural fidelity and biological activity of printed mycelium inks. Sawdust-, MEA-, and F127-based mycelium inks were used to print 3D structures and incubated for 7 days (overgrown mycelium was gently removed from the MEA-based printed structure). (**D**) Apparent viscosity under steady shear conditions was measured for sawdust-, MEA-, and F127-based mycelium inks. An uninoculated MEA-based ink was included as a control. (**E**) Storage modulus (*G*′) and loss modulus (*G*″) of four inks as a function of oscillatory strain. The crossover point of *G*′ and *G*″ indicates the yield point, marking the transition from elastic to viscous behavior. (**F**) Rheological response of mycelium inks under simulated printing conditions. The inks were subjected to alternating oscillatory (grey regions) and steady-shear (white regions) conditions, revealing their structural recovery and flow behavior. (**G**) Schematic illustrating mycelium bridging across printed filaments using mycelium ink. Hyphal growth can span certain air gaps, forming a connected network. (**H**) Bridging behavior of sawdust-based mycelium ink after incubation. No bridging indicates no hyphal growth across the gap; partial bridging shows hyphae extending without forming a full connection; full bridging refers to a continuous hyphal link between adjacent filaments. (**I**) Growth time required for the mycelium to bridge across different air gap distances (*n* = 5). (**J** and **K**) Growth behavior of *G. sessile* on MEA plates with varying malt extract concentrations after 5 days of incubation (*n*L=L5). Each plate was inoculated with an agar plug covered by actively growing mycelium (J). Radial growth radius and mean pixel intensity from 300Lpx ×L300Lpx sample regions (K).

We next examined fungal bridging across air gaps between printed filaments (**Fig. 2G** and movie S3), because mycelial bridging determines whether discrete printed filaments can grow into a continuous 3D living network. This is critical for our system, where fungal connectivity enables bacterial transport and distributed mineralization throughout the scaffold. Both *T. versicolor* and *G. sessile* fully bridged gaps up to ∼2 mm, forming continuous hyphal connections (**Figs. 2H and 2I** and fig. S3). Bridging became slower and less reliable as spacing increased, defining practical design limits for scaffold fabrication. Larger filament spacing may improve printing space and architectural openness, but it disrupts mycelial continuity and limits uniform bacterial colonization and bulk reinforcement. Therefore, scaffold spacing needs to be designed within biological bridging limits rather than based only on printing resolution. We further evaluated fungal growth on MEA plates with different malt extract concentrations to optimize 3D scaffold colonization (**Figs. 2J and 2K**). A balanced level (6% MEA) optimized both radial expansion and dense hyphal networks, while lower nutrient levels reduced density, and excessive nutrients slowed expansion (*39*, *40*). These results suggest that effective scaffold colonization requires both sufficient long-range extension and internal network density, which together support later structural continuity and biomineralization.

### Biomass-to-mineral transformation

Biomass-to-mineral transformation requires synchronizing fungal growth with bacterial activity. We therefore first allowed fungal growth to establish a continuous hyphal network throughout the printed scaffold, followed by co-culture with *S. pasteurii* in calcium- and urea-containing medium to initiate MICP. This sequence redirected mineralization from random precipitation in the surrounding solution toward localized deposition within the scaffold interior, where the mechanical framework already existed. Because effective biomineralization depends on bacterial metabolism rather than bacterial presence alone, co-culture timing was further synchronized with bacterial urease activity. Growth of *S. pasteurii* was monitored by OD_600_ (optical density at 600 nm) over 10 days (**Fig. 3A**), revealing peak activity between days 3 and 6 (OD_600_ ≈ 0.6-0.8). Co-culture was initiated during this window to maximize urease-driven carbonate generation and mineral yield. This establishes a sequential fabrication logic: fungal growth defines where mineralization can occur, while bacterial metabolism determines when reinforcement is most effective. After defining this co-culture window, we next examined whether mineralization remained surface-limited or extended throughout the scaffold volume. Micro-computed tomography (microCT) revealed relatively uniform attenuation contrast in unmineralized scaffolds but distinct high-attenuation regions within mineralized scaffolds (**Fig. 3B**), indicating dense mineral accumulation across the porous network. These internal high-density regions suggest that mineralization extended into the scaffold interior rather than remaining limited to external surfaces.

**Fig. 3.**
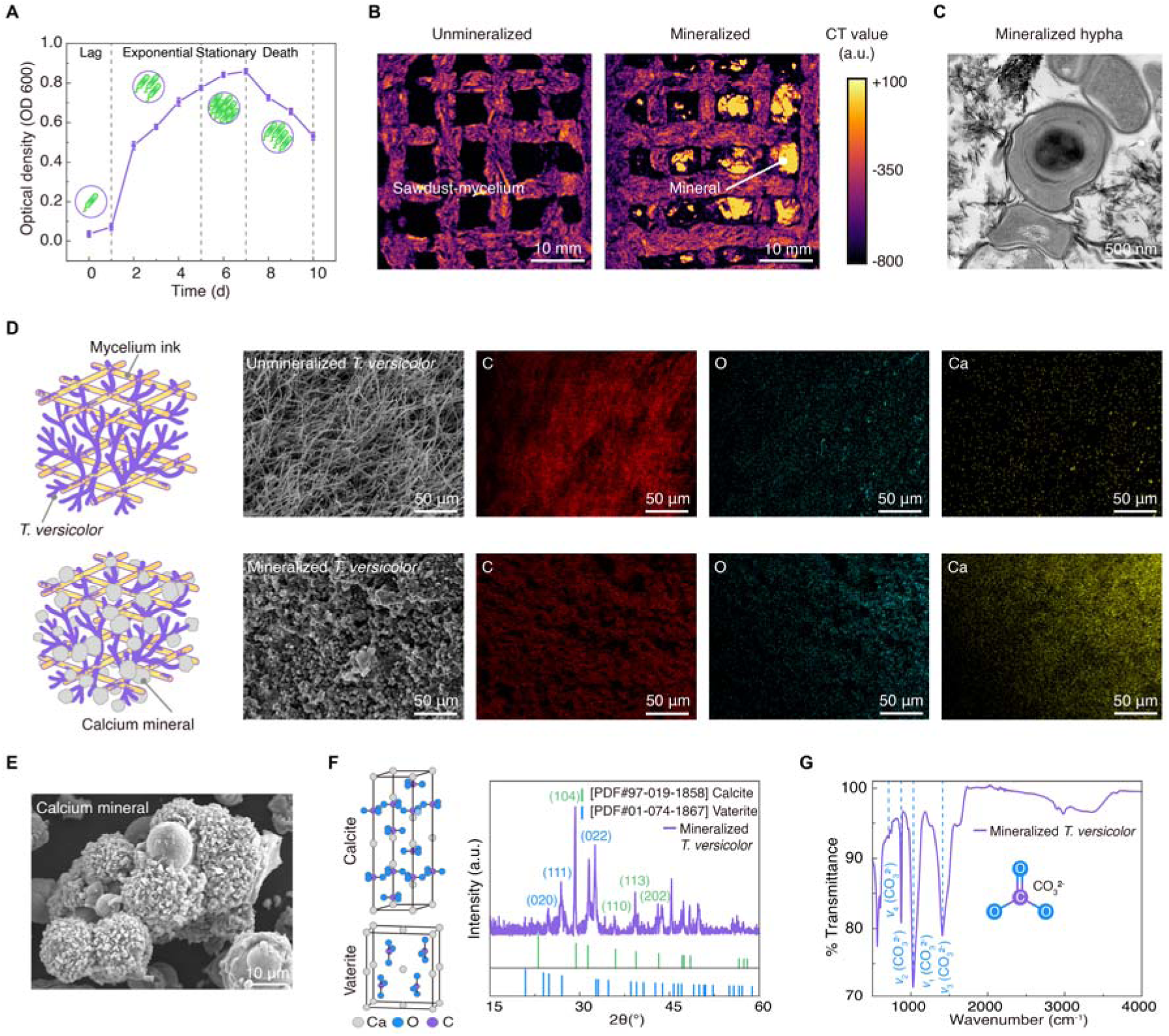
Characterization of biomineralization on mycelium scaffolds. (**A**) Growth curve of *S. pasteurii* in liquid culture measured by optical density (OD600), showing lag, exponential, stationary, and death phases over 10 days. (**B**) MicroCT images of sawdust-based mycelium scaffolds before and after mineralization. Left: The unmineralized scaffold shows a porous grid-like structure. Right: The mineralized sample displays dense white regions corresponding to calcium carbonate deposits. The grayscale bar indicates CT value (a.u.), where brighter intensities reflect higher X-ray attenuation due to mineral accumulation. (**C**) TEM image of a mineralized hypha in transverse section, showing a central lumen encircled by dark crystallites. (D) Comparison of surface structure and elemental distribution between unmineralized and mineralized *T. versicolor* mycelium scaffolds. SEM images and EDS elemental maps (C, O, Ca) reveal morphological changes and calcium carbonate deposition after biomineralization. (E) SEM image of a calcium carbonate layer collected from the surface of a *T. versicolor* mycelium scaffold mineralized through *S. pasteurii*-induced biomineralization. (**F**) XRD pattern of mineralized *T. versicolor* scaffold. Peaks are assigned to calcite and vaterite phases based on reference diffraction patterns (PDF cards). Dominant diffraction peaks at 2θL=L29.4°, 35.9°, 39.4°, and 43.2° were assigned to calcite, while additional peaks at 2θL=L24.9°, 26.9°, and 32.7° corresponded to vaterite. (**G**) FTIR spectrum of mineralized *T. versicolor*, displaying characteristic vibrational bands of carbonate ion (CO_3_^2-^). Mineralized scaffolds displayed four prominent absorption bands at 1028 cm^-1^ (ν_1_, symmetric CO_3_^2-^stretching), 873 cm^-1^ (ν_2_, out-of-plane bending), 1407 cm^-1^ (ν_3_, asymmetric stretching), and 712 cm^-1^ (ν_4_, in-plane bending).

We then examined mineralization at the microscale through transmission electron microscopy (TEM). Mineralized hyphae exhibited an electron-dense layer surrounding the fungal wall, while unmineralized hyphae retained intact cell walls and cytoplasm without obvious mineral accumulation (**Fig. 3C** and fig. S4). Mineral aggregates were also observed in adjacent extracellular regions. These observations suggest that fungal hyphae provide favorable local environments for mineral formation (*41*, *42*). Functional groups and organic residues on the fungal cell wall may support crystal nucleation, while bacterial urease activity generates local supersaturation that drives mineral growth (*43*). To further examine mineralization from microscale deposition to bulk morphology, SEM was used to compare unmineralized and mineralized scaffolds. In *T. versicolor* scaffolds, the porous hyphal network became covered by dense particulate structures, and cross-sectional images showed that mineralization extended throughout the scaffold interior rather than remaining on the surface (fig. S5). Energy-dispersive X-ray spectroscopy (EDS) further confirmed calcium-rich domains in these regions, supporting internal mineral deposition (**Fig. 3D** and fig. S6). Powder X-ray diffraction (PXRD) identified the coexistence of calcite and vaterite, a metastable polymorph commonly formed during biologically mediated mineralization (**Figs. 3E and 3F**, and fig. S7). The coexistence of calcite and vaterite suggests a biologically mediated mineralization pathway rather than simple abiotic precipitation, with rapid nucleation followed by partial maturation toward the more stable calcite phase (*44*, *45*). Fourier-transform infrared spectroscopy (FTIR) further supported carbonate formation, with characteristic absorption bands at 1028, 873, 1407, and 712 cm^-1^ observed only in mineralized scaffolds (**Fig. 3G**) (*44*). Similar mineral phases were also observed in *G. sessile* scaffolds (figs. S8-11), indicating that this process is not species-specific but arises from the general interaction between fungal hyphal networks and mineralizing bacteria. Together, these results show that biomineralization transforms porous biomass into an internal organic–inorganic composite rather than a simple surface coating.

### Bouligand living materials design for reinforcement

*In situ* biomineralization strengthens the mycelial network through local calcium carbonate deposition, but bulk structural performance depends on how effectively these reinforced regions transfer load across the entire scaffold. We then coupled biomineralization with architectural design to convert local reinforcement into macroscale load-bearing performance. Finite element analysis (FEA) of confocal-reconstructed mycelial networks showed that calcium carbonate deposition formed rigid bridges between adjacent hyphae, improving network connectivity and reducing local stress concentration (**Fig. 4A** and fig. S12). This indicates that strengthening mainly arises from enhanced load transfer between neighboring hyphae rather than simple mineral accumulation.

**Fig. 4.**
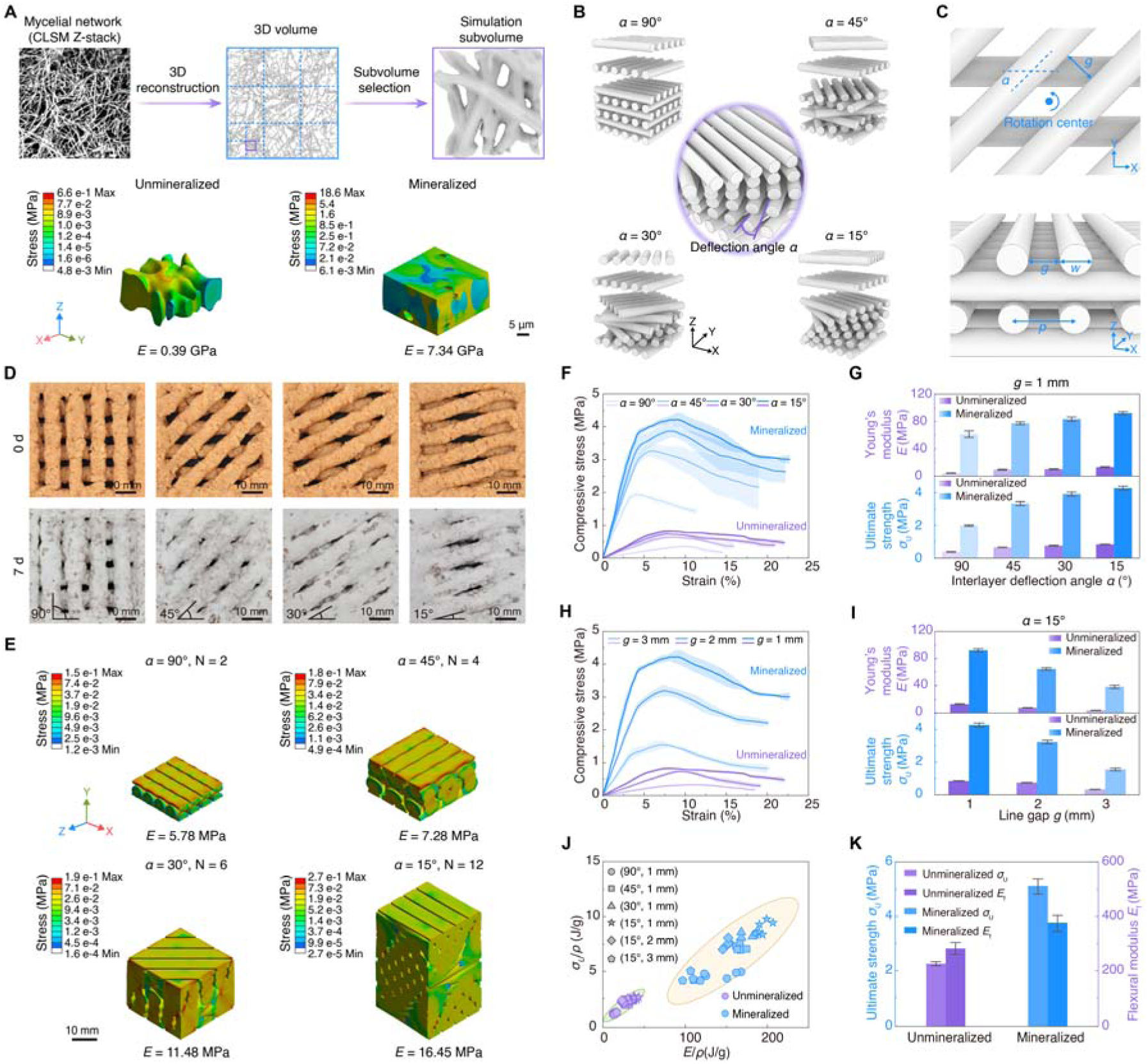
Characterization and modeling of the mechanical performance of living organic-organic composites made through the biosynthetic additive manufacturing process. (**A**) Finite element simulation of unmineralized and mineralized microscale mycelial networks. 3D geometries were reconstructed from CLSM z-stacks, followed by subvolume selection for mechanical simulation. The simulation illustrates stress distribution and computes the effective Young’s modulus of the networks. Mineralization was modeled by partially filling hyphal gaps with calcium carbonate. (**B**) Design of multilayered helicoidal scaffolds with varying interlayer deflection angles (α = 90°, 45°, 30°, 15°), forming a Bouligand structure. (**C**) Schematic diagrams of the designed Bouligand scaffold. Top: Interlayer deflection angle (α) defines the rotational offset between adjacent layers. Bottom: Printing path parameters including interline gap (*g*), line width (*w*), and center-to-center spacing (*p*) between adjacent lines. (**D**) Top-view images of printed scaffolds with varying interlayer deflection angles, showing adjacent-layer geometry. Images were captured at day 0 and after 7 days of fungal growth. (**E**) Finite element simulation showing von Mises stress distribution under uniaxial compression for scaffolds with different interlayer deflection angles (α = 90°, 45°, 30°, and 15°). Stress concentration patterns vary with angle. (**F** and **G**) Compressive mechanical performance of unmineralized (purple) and mineralized (blue) mycelium scaffolds with varying interlayer deflection angles (α = 90°, 45°, 30°, and 15°; *g* = 1 mm). Compressive stress-strain curves with dark lines as interpolated means, light shading as mean ± SD (upper = mean + SD, lower = mean - SD) (F). Young’s modulus and ultimate strength (G) compare unmineralized and mineralized conditions (*n* = 5). (**H** and **I**) Compressive mechanical performance of unmineralized (purple) and mineralized (blue) mycelium scaffolds with different line gaps (*g* = 1, 2, and 3 mm; α = 15°). Compressive stress-strain curves with dark lines as interpolated means, light shading as mean ± SD (upper = mean + SD, lower = mean - SD) (H). Young’s modulus and ultimate strength (I), comparing unmineralized and mineralized conditions (*n* = 5). (**J**) Ashby plots of specific ultimate strength and specific Young’s modulus (normalized by bulk density) for mineralized and unmineralized composites with different structural configurations. The 95% confidence ellipses are shown to indicate the statistical spread and central tendency of each group. (**K**) Flexural modulus and ultimate flexural strength of unmineralized (purple) and mineralized (blue) specimens measured from three-point bending tests (*n* = 5).

To further translate this microscale reinforcement into bulk mechanics, we designed multilayer Bouligand scaffolds by varying interlayer deflection angle (α = 90°, 45°, 30°, and 15°) and interline gap (*g* = 1, 2, and 3 mm) (**Figs. 4B-D** and fig. S13). These two parameters were chosen because α controls stress transfer between adjacent layers, while *g* determines whether fungal hyphae can bridge neighboring filaments and support continuous bacterial colonization and internal mineralization (*46*). Smaller α values increased filament overlap and reduced stress concentration, leading to more continuous load transfer (**Fig. 4E** and fig. S14). Compression tests further confirmed that decreasing α from 90° to 15° significantly improved stiffness and strength in both unmineralized and mineralized scaffolds (**Figs. 4F and 4G**). In mineralized samples, Young’s modulus increased from 61.23 ± 7.09 MPa to 92.18 ± 3.28 MPa, while compressive strength increased from 1.98 ± 0.06 MPa to 4.26 ± 0.18 MPa (table S1). Relative to their corresponding unmineralized controls, mineralization improved modulus by 7-14-fold and strength by over 5-fold, demonstrating strong synergy between Bouligand architecture and calcium carbonate deposition. When α was fixed at 15°, the strongest reinforcement was achieved at 1 mm spacing (**Figs. 4H and 4I**, and table S2), where dense fungal networks enabled efficient hyphal bridging, load transfer, and internal mineralization.

Ashby plots further showed that mineralized scaffolds shifted toward higher specific strength (σ_U_ /ρ) and modulus (*E*/ρ), confirming that reinforcement came from structural optimization rather than simple densification (**Fig. 4J**). Control experiments verified that this transition required both fungal growth and bacterial mineralization: sawdust-only scaffolds collapsed in mineralization medium, while dry scaffolds without living mycelium showed much weaker performance (figs. S15 and S16, and table S3). Compression and bending exhibited distinct responses to mineralization due to different stress distributions (*47*). Compression mainly benefits from internal mineralization, while bending is less affected because mineral deposition is concentrated inside the scaffold rather than at the surface (figs. S17-20 and movies S4-5). As a result, compressive performance improved substantially, while flexural modulus increased only moderately from 282.43 MPa to 373.99 MPa (**Fig. 4K** and table S4). This confirms that internal volumetric mineralization is the dominant strengthening mechanism for compressive load-bearing applications.

### Self-healing and modular assembly

Beyond mineral-enabled strengthening, the living nature of microbial networks enables autonomous self-healing to restore structural continuity, and modular self-assembly to form larger complex structures (*48*, *49*). As shown in **Fig. 5A**, artificially scratched scaffolds regained continuity after 7 days of hyphal regrowth. This behavior addresses one of the fundamental limitations of extrusion-based 3D printing, where surface cracks, weak interlayer interfaces, and local discontinuities are often unavoidable during deposition (*50*, *51*). Instead of requiring external repair, living mycelium can regrow across damaged regions and restore structural continuity through active hyphal extension (movie S6). This allows the scaffold to recover after fabrication rather than relying entirely on highly precise extrusion during printing. Mechanical testing confirmed that both self-healed and self-assembled scaffolds recovered compressive modulus and strength, with mineralized samples showing further enhancement compared to unmineralized ones (**Figs. 5B and 5C**, and table S5). This demonstrates that healing is not only visual reconnection, but also functional structural recovery. After bacterial biomineralization, repaired regions regained stronger load-bearing capacity, indicating that biological healing and mineral reinforcement act sequentially to restore and strengthen damaged architectures.

**Fig. 5.**
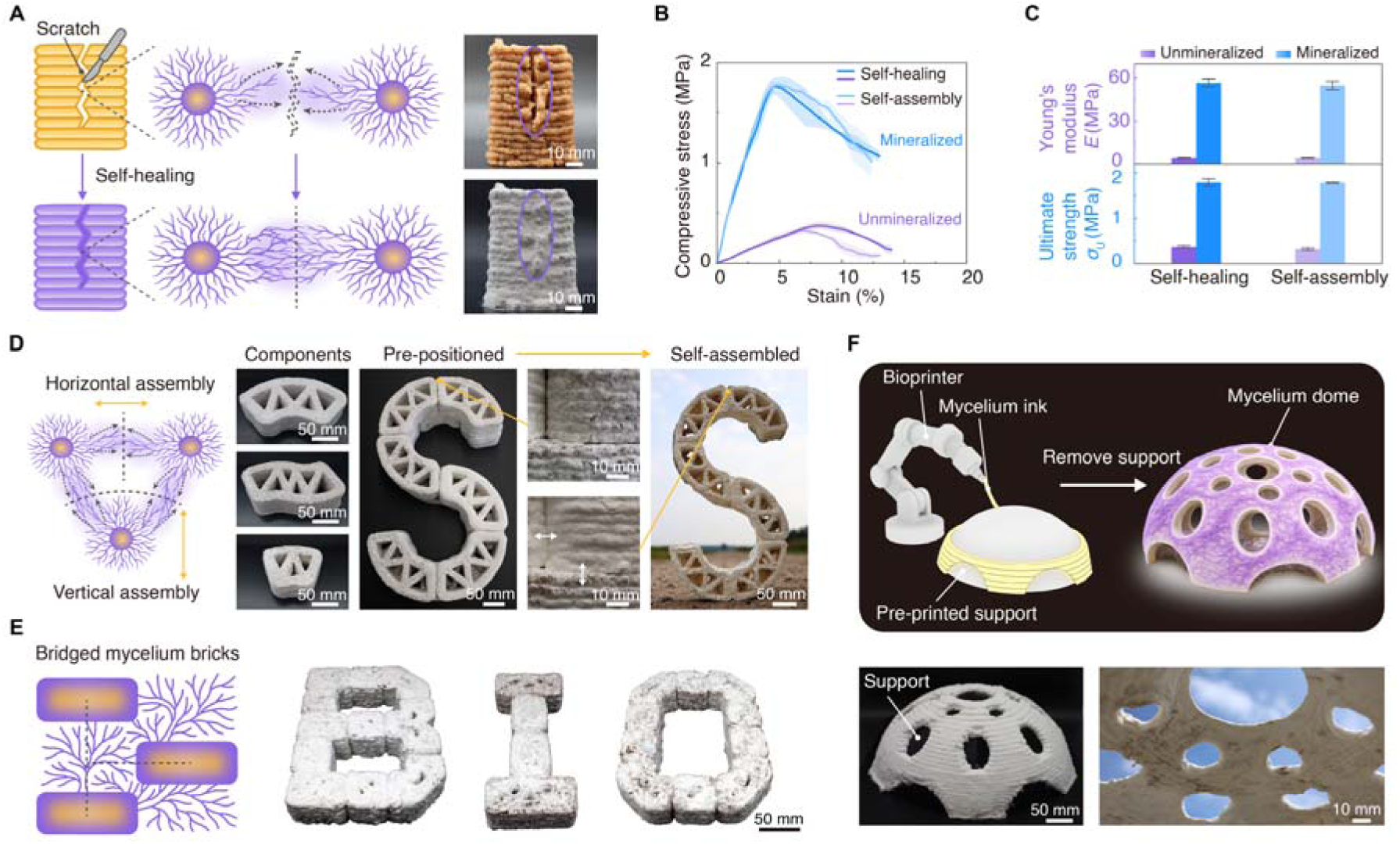
Self-healing and modular self-assembly capacities of the 3D-printed living structures. (**A**) Self-healing mycelium scaffold. A printed scaffold with an introduced scratch is gradually healed through hyphal growth, resulting in the restoration of an integrated structure. (**B** and **C**) Compressive mechanical performance of self-healing and self-assembly mycelium scaffolds (*g* = 1 mm; α = 90°). Compressive stress-strain curves with dark lines as interpolated means, light shading as mean ± SD (upper = mean + SD, lower = mean - SD) (B). Young’s modulus and ultimate strength (C), comparing unmineralized (purple) and mineralized (blue) conditions (*n* = 3). (**D** and **E**) Demonstration of self-assembly behaviors in printed mycelium modules. An “S”-shaped pattern formed from pre-positioned bricks of varying sizes assembled by hyphal adhesion (D). The letters “BIO” are printed as discrete modules and connected through hyphal bridging across air gaps (E). (**F**) A dome structure fabricated by printing mycelium ink over a removable support, resulting in a freestanding hollow architecture after support removal.

Instead of repairing damage within a single scaffold, we showed that multiple independently bioprinted living units can be positioned separately and then fused into one continuous structure through biological growth. This is the central rationale for modular fabrication: large or complex structures are difficult to print as a single piece, but smaller inoculated modules can be fabricated efficiently, transported easily, and assembled on-site. Through subsequent hyphal growth, adjacent modules gradually become biologically bonded, as structures are perhaps too large or too complex to be printed directly in 3D. In this way, mycelium functions as a self-growing biological binder that strengthens the connection over time and enables seamless structural integration after assembly. Pre-positioned components were arranged with controlled gaps, where hyphal growth bridged adjacent interfaces to form stable joints (**Figs. 5D and 5E**, and figs. S21-22). Initial placement provided geometric alignment, while mycelial growth created long-term fusion and reinforcement. These results show that structural continuity can emerge after assembly rather than being fully predefined during printing, which is essential for scalable living construction materials.

Because the bioink must maintain sufficient moisture to support biological growth, large-span or hollow overhanging geometries remain difficult to fabricate directly (fig. S23 and movie S7). To address this limitation, we deposited mycelium ink onto a pre-printed removable polylactic acid (PLA) support to fabricate a dome-shaped structure (**Fig. 5F** and movie S8). After the removal of the support, the dome remained standing, demonstrating a practical strategy for constructing suspended architectures using living bioinks. Here, the temporary synthetic support provides short-term printability, while the living scaffold gradually becomes self-supporting through mycelial growth. Together, these results establish a biosynthetic additive manufacturing framework in which digital fabrication and biological growth are integrated to produce self-healing, self-assembling, and scalable structural materials.

## DISCUSSION

This work establishes biosynthetic additive manufacturing as a unique strategy to convert lignocellulosic biomass into mechanically robust living structural composites through sequential biological activity. Rather than serving merely as a biological filler, mycelium functions as the structural framework that enables bacteria to move beyond surface mineralization and achieve volumetric reinforcement throughout the scaffold. Mechanical strengthening emerges from the synergistic coupling of biomineralization and Bouligand architecture, where geometric organization governs how effectively mineralized junctions contribute to bulk load transfer and structural integrity. Beyond mechanical performance, the living microbial network introduces functionalities absent from conventional synthetic materials, including self-healing to restore structural continuity and modular self-assembly into complex architectures, thereby providing a scalable and defect-tolerant strategy for next-generation construction materials.

More broadly, this work introduces a new paradigm for engineered living systems by integrating microbial consortia with inert structural components in a synergistic physical-biological platform inspired by natural ecological systems. Fungal-bacterial interactions and microbial interdependencies are harnessed to regulate habitation, reproduction, and long-term material evolution across multiple time and length scales. Simultaneously, the study advances materials science and additive manufacturing through a new biomanufacturing framework that seamlessly combines architected 3D printing with biosynthetic growth processes for programmable materials design using scalable and sustainable biomass feedstocks. From an environmental perspective, the platform avoids energy-intensive thermal processing by relying on microbial mineralization, offering a potential pathway toward carbon-negative construction. The integration of renewable biomass utilization, self-healing capability, modular reuse, and extended service lifetime may further reduce lifecycle emissions compared with conventional cement-based materials, while establishing new opportunities to expand the manufacturing capacity and functionality of hybrid, living, manmade systems.

## MATERIALS AND METHODS

Detailed experimental procedures, instrument settings, and data-processing methods are provided in the Supplementary Materials.

### Materials

Soybean peptone (CAS 73049-73-7), Pluronic F127 (F127, CAS 9003-11-6), tryptone (CAS 91079-40-2), yeast extract (CAS 8013-01-2), calcofluor white (CFW, CAS 18909-50-5), urea (CO(NH_2_)_2_, CAS 57-13-6), sodium chloride (NaCl, CAS 7647-14-5), ammonium chloride (NH_4_Cl, CAS 12125-02-9), sodium phosphate dibasic (Na_2_HPO_4_, CAS 7558-79-4), formaldehyde (CH_2_O, 37 wt%, CAS 50-00-0), and phosphate-buffered saline (PBS, pH 7.2) were purchased from Sigma-Aldrich, USA. Malt extract (CAS 8002-48-0) and κ-carrageenan (CAS 11114-20-8) were purchased from Research Products International, USA. Agar (CAS 9002-18-0) and calcium chloride dihydrate (CaCl_2_·2H_2_O, CAS 10035-04-8) were purchased from Carolina Biological Supply Co., USA. Sodium hydroxide (NaOH, 98.0%, CAS 1310-73-2), hydrochloric acid (HCl, 35 wt%, CAS 7647-01-0), and Alizarin Red S sodium salt (ARS, 1% w/v, CAS 130-22-3) were purchased from Thermo Fisher Scientific, USA. Microcrystalline cellulose (MCC, CAS 9004-34-6, Loudwolf Industrial & Scientific, USA), carboxymethyl cellulose (CMC, CAS 9000-11-7, La Tiendita Essentials, USA), and propidium iodide (PI, CAS 25535-16-4, Life Technologies, USA) were used as received. Glutaraldehyde (25% aqueous solution, CAS 111-30-8), paraformaldehyde (16% aqueous solution, CAS 30525-89-4), osmium tetroxide (CAS 20816-12-0), uranyl acetate (CAS 6159-44-0), acetone (CAS 67-64-1) and Spurr resin (CAS 52368-54-4) were purchased from Electron Microscopy Sciences, USA. *Trametes versicolor* (*T. versicolor*) sawdust spawn and *Ganoderma sessile* (*G. sessile*) were obtained from Field & Forest Products, USA. *Sporosarcina pasteurii* (*S. pasteurii*, ATCC #11859) was obtained from the American Type Culture Collection, USA. Deionized (DI) water used for all solution preparation and sample processing was obtained from a Thermo Scientific water purification system (Thermo Fisher Scientific Inc., USA).

### Preparation of mycelium inks

F127-based mycelium ink was prepared by dissolving 30% (w/v) F127 in pre-chilled 2% (w/v) malt extract medium containing 0.5% (w/v) peptone at pH 6.8. The solution was sterilized using a 0.22-μm PES filter. *G. sessile* was cultured in the same medium for 5 days, collected by centrifugation at 4000 rpm for 10 min, and dispersed in 50 mL of sterile F127 solution. The resulting ink was stored at 4 °C and gently mixed before use.

MEA-based ink contained 6% (w/v) malt extract, 4.5% (w/v) agar, 1.5% (w/v) κ-carrageenan, 0.5% (w/v) peptone, 1% (w/v) MCC, and 1% (w/v) CMC. The formulation was homogenized and autoclaved at 121 °C for 20 min. After solidification and mechanical disruption, the ink was inoculated with a 1 cm × 1 cm *G. sessile* agar plug and incubated at 28 °C, 95% relative humidity (RH), and in the dark for 7 to 10 days. The surface mycelium and inoculation plug were removed, and the remaining ink was rehomogenized before printing.

Sawdust-based ink was prepared using the same formulation without malt extract and peptone. The sterilized matrix was combined with 25% (w/w) *T. versicolor* sawdust spawn and homogenized at 1600 rpm for 2 min using 15-mm grinding media. Dense surface mycelium was removed before mixing to prevent nozzle blockage. An autoclaved mycelium-containing sawdust formulation was prepared using the same procedure as a control.

### Rheology of mycelium inks

The rheological properties of F127-based mycelium ink, MEA-based pure ink, MEA-based mycelium ink, and sawdust-based mycelium ink were measured using a Discovery HR-2 rheometer with a 40-mm parallel plate and a 1000-μm gap at 25 °C. Apparent viscosity was measured over shear rates of 0.1 to 100 s^-1^. Oscillatory strain sweeps were conducted from 0.1 to 100% strain at 10 rad s^-1^ to determine the storage modulus, loss modulus, and yield point. Structural recovery was evaluated through repeated cycles of oscillation at 1% strain and 10 rad s^-1^ for 60 s, followed by steady shear at 1 s^-1^ for 30 s.

### Preparation of growth media and fungal culture

*G. sessile* was maintained on MEA containing 2.0% (w/v) malt extract, 0.5% (w/v) peptone, and 1.5% (w/v) agar at pH 6.8. The medium was autoclaved at 121 °C for 20 min. A 1 cm × 1 cm mycelium-covered agar plug was transferred to fresh MEA every four weeks and incubated at 28 °C, 95% RH, and in the dark.

Liquid cultures were prepared using 2.0% (w/v) malt extract and 0.5% (w/v) peptone at pH 6.8. Mycelium-covered agar plugs were transferred into 50 mL of liquid medium and cultured under the same temperature and light conditions. Commercially prepared, fully colonized *T. versicolor* hardwood sawdust spawn was stored at 4 °C until use.

### Mycelial growth behaviour and bridging across air gaps

The effects of nutrient concentration on *G. sessile* growth were evaluated on MEA containing 0 to 10% (w/v) malt extract. Plates were inoculated with 1 cm × 1 cm mycelium-covered plugs and incubated at 28 °C, 95% RH, and in the dark for 5 days. Radial expansion and mean pixel intensity within 300 pixel × 300 pixel regions were quantified using ImageJ. Five regions were measured per plate, with five replicate plates per condition.

Mycelial bridging was assessed by printing parallel filaments separated by controlled air gaps. Sawdust-based *T. versicolor* ink was printed using a 6-mm nozzle with gaps of 1 to 7 mm, whereas MEA-based *G. sessile* ink was printed using a 1.6-mm nozzle with gaps of 0.5 to 2.5 mm. Samples were incubated at 28 °C and 95% RH for 7 days for *T. versicolor* or 4 days for *G. sessile*. Bridging was defined as visible hyphal growth connecting most of the gap between adjacent filaments.

## 3D printing of mycelium-based mechanical specimens

Mechanical specimens were printed using a 3D PotterBot 10 PRO at 20 mm s^-1^. Models were designed in SolidWorks and converted to printing paths using UltiMaker Cura. Sawdust-based mycelium ink was extruded through a 6-mm nozzle onto sterilized polyvinyl chloride sheets.

Compression specimens consisted of 16 printed layers enclosed by a single-shell outer wall. Adjacent layers were rotated by interlayer deflection angles of 15°, 30°, 45°, or 90°, with interline gaps of 1, 2, or 3 mm. Three-point bending specimens consisted of four layers with 90° filament orientation and a 1-mm gap. Printed specimens were incubated at 28 °C, 95% RH, and in the dark for 7 days and subsequently dried at 70 °C to constant weight.

### Mechanical compression and three-point bending tests

Uniaxial compression tests were performed at 25 °C using a universal testing machine equipped with a 20,000-lb load cell. Five specimens were tested per condition. Sample dimensions were measured at five positions and averaged to calculate the initial height and cross-sectional area. Specimens were compressed at 6 mm min^-1^ until failure. Compressive strain and stress were calculated from displacement, initial height, applied force, and cross-sectional area. Young’s modulus was obtained by linear fitting over the 0 to 1% strain region, and ultimate compressive strength was defined as the maximum stress before failure. Bulk density was calculated from dry mass and geometric volume.

Three-point bending was conducted using a 100-lb load cell, a 57-mm support span, and a loading rate of 6 mm min^-1^ at 25 °C. Flexural stress and strain were calculated from specimen dimensions, applied force, support span, and midspan deflection. Flexural modulus was determined from the initial 0 to 0.5% strain region, and flexural strength was defined as the maximum stress before failure. Data were analyzed in Origin 2021. Representative stress–strain curves were generated by interpolation at 0.02% strain intervals across the common strain range.

### Bacteria culturing and biomineralization of mycelium scaffolds

*S. pasteurii* was cultured in alkaline peptone yeast extract medium containing 1.0% (w/v) tryptone, 0.5% (w/v) yeast extract, 0.5% (w/v) NaCl, 0.5% (w/v) NH_4_Cl, and 0.1% (w/v) Na_2_HPO_4_ at pH 8.0. Cultures were incubated at 28 °C and 100 rpm. Bacterial growth was monitored daily by OD600 measurements, with five measurements averaged at each time point.

Cultures were harvested at the late exponential phase at OD600 values of approximately 0.6 to 0.8, centrifuged at 4000 rpm for 10 min, and resuspended in fresh medium. Mycelium scaffolds pre-incubated for 7 days were completely submerged in the bacterial suspension at a liquid-to-solid volume ratio greater than 2:1 and cultured at 28 °C and 100 rpm for 24 hours.

Biomineralization was initiated by replacing half of the bacterial suspension with an equal volume of calcium-based mineralization medium containing 2.0% (w/v) urea, 1.0% (w/v) CaCl_2_·2H_2_O, 0.1% (w/v) yeast extract, and 0.1% (w/v) Na_2_HPO_4_ at pH 8.0. Urea was filter-sterilized separately and added after autoclaving the remaining components. Scaffolds were incubated for an additional 48 hours, removed from the medium, and dried at 70°C to constant weight.

### Preparation of self-healing and self-assembly mycelial scaffolds

Self-healing and self-assembly specimens were fabricated using the same printing, incubation, biomineralization, drying, and compression conditions as the other compression specimens, with an interlayer angle of 90° and a 1-mm line gap. For self-healing experiments, a sterile scalpel was used to introduce a surface scratch approximately five filament layers in length immediately after printing. Samples were then incubated to allow hyphal regrowth before drying and testing.

For self-assembly experiments, scaffolds were printed as two equal-height sections. The sections were incubated separately for 7 days, manually placed together, and incubated for an additional 7 days to allow hyphal fusion before drying and testing.

## 3D printing of mycelium inks and self-assembly of printed structures

F127- and MEA-based inks were printed using a Brinter ONE bioprinter equipped with a pneumatic extrusion head and 18-gauge and 14-gauge tips, respectively. Printing was performed at 6 mm s^-1^ onto sterile glass substrates. F127 ink was loaded into sterile cartridges at 4 °C and equilibrated at room temperature for 5 min before printing.

Sawdust-based inks were printed under the same general conditions used for mechanical specimens. Overhanging structures were deposited onto removable polylactic acid supports fabricated using fused deposition modeling. The supports were removed after mycelial growth and drying.

For modular assembly, individual printed components were incubated at 28 °C and 95% RH for 7 days, positioned adjacent to one another or held in a predefined arrangement, and incubated for another 7 days. Hyphal extension across neighboring interfaces connected the separate modules without external adhesives.

### Confocal laser scanning microscopy of co-cultured mycelium and bacteria

Fragments of *T. versicolor* or *G. sessile* were co-cultured with late-exponential-phase *S. pasteurii* at 28 °C and 100 rpm for 24 hours. Samples were fixed with 4% (w/v) formaldehyde in 0.1 M PBS for 20 min and washed twice. Fungal hyphae and bacterial cells were stained with 0.25 g L^-1^ CFW and 0.1 g L^-1^ PI, respectively, for 15 min in the dark.

Mineralized samples were stained with CFW and 1% (w/v) ARS for 15 min without subsequent washing to preserve the hydrated mineral phase. Samples were imaged using a Nikon Eclipse Ni-U microscope equipped with a C2+ confocal system and a 20×/0.75 numerical aperture objective. CFW was excited at 405 nm, whereas PI and ARS were excited at 561 nm. Z-stacks were collected at 1-μm intervals and rendered using NIS-Elements. Only uniform linear brightness and contrast adjustments were applied.

### Scanning electron microscopy and energy dispersive X-ray microanalysis

Dried unmineralized and mineralized scaffolds were mounted on aluminum stubs and coated with approximately 10 nm of osmium. Cross sections were prepared using a low-speed diamond saw. Samples were imaged using a JEOL JSM-6610LV scanning electron microscope at an accelerating voltage of 12 kV with a secondary electron detector. Elemental analysis and mapping were performed using an Oxford Instruments AZtec energy-dispersive X-ray spectroscopy system equipped with a 20-mm^2^ silicon drift detector.

### Powder X-ray diffraction and Fourier-transform infrared spectroscopy

Dried scaffolds were ground into homogeneous powders. Powder X-ray diffraction patterns were collected using Cu-Kα_1_ radiation at 40 kV and 40 mA over a 2θ range of 15° to 60°, with a step size of 0.02° and a counting time of 0.05 s per step. Mineral phases were identified using JADE 9 software.

Fourier-transform infrared spectra were acquired from the same powdered samples using a Spectrum Two spectrometer equipped with an attenuated total reflectance accessory. Spectra were recorded from 4000 to 500 cm^-1^ at a resolution of 4 cm^-1^ and averaged over 32 scans.

### Micro-computed tomography

Dry mineralized and unmineralized *T. versicolor* scaffolds were scanned using a Quantum GX micro-computed tomography system at 90 kV and 88 μA with a 72 mm × 72 mm field of view. Images were acquired with a voxel size of 144 μm and an 18-s scan duration. Samples were reconstructed at the same resolution. A uniform display window with a center of −350 arbitrary units and a width of 900 arbitrary units was applied to both conditions. Representative slices were converted to 8-bit images and pseudo-colored using the mpl-inferno lookup table in Fiji.

### Transmission electron microscopy

Samples were fixed in 2.5% (w/v) glutaraldehyde and 2.0% (w/v) paraformaldehyde in 0.1 M PBS, rinsed, and post-fixed with 1% (w/v) osmium tetroxide. Samples were stained with 1% (w/v) uranyl acetate, dehydrated through a graded acetone series, and embedded in Spurr resin. Ultrathin 70-nm sections were prepared, post-stained with uranyl acetate and lead citrate, and imaged using a JEOL 1400Flash transmission electron microscope at 100 kV.

### Compression simulation of the Bouligand structure

Bouligand scaffolds were modeled in SolidWorks and analyzed using ANSYS Workbench. Printed filaments were represented as solid cylinders with diameters matching the extrusion nozzle, a 1-mm line gap, and controlled overlap between adjacent layers. Overlapping regions were treated as bonded. Structures with interlayer deflection angles of 15°, 30°, 45°, and 90° were subjected to a uniaxial strain of 0.0015 under linear elastic conditions. Effective Young’s modulus was calculated from the simulated stress–strain response.

### Compression simulation of microscale unmineralized and mineralized mycelial networks

Three-dimensional mycelial geometries were reconstructed from CLSM z-stacks of *T. versicolor* using Fiji and converted to volumetric models using Rhino 8. A representative 24 μm × 24 μm × 15 μm subvolume was selected based on typical hyphal density and connectivity. The unmineralized model contained only mycelial filaments, whereas the mineralized model included calcium carbonate within the pore regions.

Mineralized porosity was set to 16.17% based on binarized microCT images and represented using randomly distributed spherical pores. Calcium carbonate and mycelium interfaces were defined as bonded. A compressive displacement of 0.001 μm was applied to the top surface while the bottom surface was fixed. The mycelial Young’s modulus and Poisson’s ratio were set to 561 MPa and 0.25, respectively, based on reported properties of dimitic basidiomycete hyphae. The corresponding values for biologically precipitated calcium carbonate were 33.98 GPa and 0.31. Material parameters were derived from previously reported nanoindentation measurements.

## Acknowledgments

**Funding:**

National Science Foundation grant ECCS-2339495 (JL)

National Science Foundation grant ECCS-2334134 (JL)

National Science Foundation grant ECCS-2216131 (JL)

National Science Foundation grant FEMA-2318057 (JL)

National Science Foundation grant CMMI-2323917 (JL)

**Author contributions:**

Conceptualization: J.L.

Data curation: H.Y., L.H., L.W.

Supervision: J.L.

Funding acquisition: J.L.

Investigation: H.Y., L.H., Y.X., L.W., K.N., Y.C., C.C., G.H., R.Y., E.D., M.B., S.M., S.I., X.Z., J.H.

Visualization: H.Y.

Resources: B.W., X.S., T.L., G.B.

Methodology: H.Y., L.H., J.H., Y.X., L.W., K.N., B.W., T.L., G.B., K.L., J.L.

Writing – original draft: H.Y., J.L.

Writing – review & editing: H.Y., J.L.

**Competing interests**

H.Y. and J.L. are inventors on a patent application (No. TEC2026-0174) submitted by the Michigan State University Innovation Center.

**Data, code, and materials availability**

All data are available in the main text or the supplementary materials.

## Supplementary information

Materials and Methods

Supplementary Text

Figs. S1-23

Table S1-5

Movies S1-8

## REFERENCES

1. G. Churkina, A. Organschi, C. P. O. Reyer, A. Ruff, K. Vinke, Z. Liu, B. K. Reck, T. E. Graedel, H. J. Schellnhuber, Buildings as a global carbon sink. Nat. Sustain. 3, 269–276 (2020).

2. A. Mishra, F. Humpenöder, G. Churkina, C. P. O. Reyer, F. Beier, B. L. Bodirsky, H. J. Schellnhuber, H. Lotze-Campen, A. Popp, Land use change and carbon emissions of a transformation to timber cities. Nat. Commun. 13, 4889 (2022).

3. A. Sturcová, G. R. Davies, S. J. Eichhorn, Elastic modulus and stress-transfer properties of tunicate cellulose whiskers. Biomacromolecules 6, 1055–1061 (2005).

4. A. Dufresne, Nanocellulose: from nature to high performance tailored materials. (2017).

5. S. R. Johnston, L. Boddy, A. J. Weightman, Bacteria in decomposing wood and their interactions with wood-decay fungi. FEMS Microbiol. Ecol. 92, fiw179 (2016).

6. J. Cai, Y. He, X. Yu, S. W. Banks, Y. Yang, X. Zhang, Y. Yu, R. Liu, A. V. Bridgwater, Review of physicochemical properties and analytical characterization of lignocellulosic biomass. Renew. Sustain. Energy Rev. 76, 309–322 (2017).

7. N. W. Sokol, E. Slessarev, G. L. Marschmann, A. Nicolas, S. J. Blazewicz, E. L. Brodie, M. K. Firestone, M. M. Foley, R. Hestrin, B. A. Hungate, B. J. Koch, B. W. Stone, M. B. Sullivan, O. Zablocki, LLNL Soil Microbiome Consortium, J. Pett-Ridge, Life and death in the soil microbiome: how ecological processes influence biogeochemistry. Nat. Rev. Microbiol. 20, 415–430 (2022).

8. P. Anbu, C.-H. Kang, Y.-J. Shin, J.-S. So, Formations of calcium carbonate minerals by bacteria and its multiple applications. Springerplus 5, 250 (2016).

9. W. Qin, C.-Y. Wang, Y.-X. Ma, M.-J. Shen, J. Li, K. Jiao, F. R. Tay, L.-N. Niu, Microbe-mediated extracellular and intracellular mineralization: Environmental, industrial, and biotechnological applications. Adv. Mater. 32, e1907833 (2020).

10. A. Spadafora, E. Perri, J. A. Mckenzie, C. Vasconcelos, Microbial biomineralization processes forming modern Ca:Mg carbonate stromatolites: Biomineralization processes forming stromatolites. Sedimentology 57, 27–40 (2010).

11. W. Gwenzi, ‘Carbon sequestration via biomineralization: Processes, applications and future directions’ in Sustainable Agriculture Reviews (Springer International Publishing, Cham, 2019), pp. 93–106.

12. M. P. Nelsen, W. A. DiMichele, S. E. Peters, C. K. Boyce, Delayed fungal evolution did not cause the Paleozoic peak in coal production. Proc. Natl. Acad. Sci. U. S. A. 113, 2442–2447 (2016).

13. F. Jiang, T. Li, Y. Li, Y. Zhang, A. Gong, J. Dai, E. Hitz, W. Luo, L. Hu, Wood-based nanotechnologies toward sustainability. Adv. Mater. 30, 1703453 (2018).

14. R. M. McBee, M. Lucht, N. Mukhitov, M. Richardson, T. Srinivasan, D. Meng, H. Chen, A. Kaufman, M. Reitman, C. Munck, D. Schaak, C. Voigt, H. H. Wang, Engineering living and regenerative fungal-bacterial biocomposite structures. Nat. Mater. 21, 471–478 (2022).

15. S. Gantenbein, E. Colucci, J. Käch, E. Trachsel, F. B. Coulter, P. A. Rühs, K. Masania, A. R. Studart, Three-dimensional printing of mycelium hydrogels into living complex materials. Nat. Mater. 22, 128–134 (2023).

16. L. Yang, Z. Qin, Mycelium-based wood composites for light weight and high strength by experiment and machine learning. Cell Rep. Phys. Sci. 4 (2023).

17. G. De, L. Yang, J. Lee, Y.-H. Wu, Z. Tian, Z. Qin, Mycelium–coir-based composites for sustainable building insulation. J. Mater. Chem. A Mater. Energy Sustain. 13, 9694–9707 (2025).

18. R. Abhijith, A. Ashok, C. R. Rejeesh, Sustainable packaging applications from mycelium to substitute polystyrene: a review. Mater. Today 5, 2139–2145 (2018).

19. S. Sivaprasad, S. K. Byju, C. Prajith, J. Shaju, C. R. Rejeesh, Development of a novel mycelium bio-composite material to substitute for polystyrene in packaging applications. Mater. Today 47, 5038–5044 (2021).

20. M. G. Pelletier, G. A. Holt, J. D. Wanjura, E. Bayer, G. McIntyre, An evaluation study of mycelium based acoustic absorbers grown on agricultural by-product substrates. Ind. Crops Prod. 51, 480–485 (2013).

21. E. S. L. Vasquez, K. Vega, Myco-Accessories: Sustainable Wearables with Biodegradable Materials (2019)Proceedings of the 2019 ACM international symposium on wearable computers.

22. B. An, Y. Wang, Y. Huang, X. Wang, Y. Liu, D. Xun, G. M. Church, Z. Dai, X. Yi, T.-C. Tang, Engineered living materials for sustainability. Chem. Rev. 123, 2349–2419 (2022).

23. A. Rodrigo-Navarro, S. Sankaran, M. J. Dalby, A. del Campo, M. Salmeron-Sanchez, Engineered living biomaterials. Nat. Rev. Mater. 6, 1175–1190 (2021).

24. N. K. Dhami, M. S. Reddy, A. Mukherjee, Biomineralization of calcium carbonates and their engineered applications: a review. Front. Microbiol. 4, 314 (2013).

25. E. Viles, E. Heyneman, S. Lin, V. Montague, A. Darabi, L. M. Cox, A. Phillips, R. Gerlach, E. J. Espinosa-Ortiz, C. Heveran, Mycelium as a scaffold for biomineralized engineered living materials. Cell Rep. Phys. Sci. 6 (2025).

26. M. J. Castro-Alonso, L. E. Montañez-Hernandez, M. A. Sanchez-Muñoz, M. R. Macias Franco, R. Narayanasamy, N. Balagurusamy, Microbially induced calcium carbonate precipitation (MICP) and its potential in bioconcrete: Microbiological and molecular concepts. Front. Mater. 6 (2019).

27. Y. Wu, H. Li, Y. Li, Biomineralization Induced by Cells of Sporosarcina pasteurii: Mechanisms, Applications and Challenges. Microorganisms 9, 2396 (2021).

28. T. S. Moon, K. Yum, Integrating Engineered Living Materials with 3D Bioprinting. Adv. Funct. Mater., 2500934 (2025).

29. S. C. Shen, N. A. Lee, W. J. Lockett, A. D. Acuil, H. B. Gazdus, B. N. Spitzer, M. J. Buehler, Robust myco-composites: a biocomposite platform for versatile hybrid-living materials. Mater. Horiz. 11, 1689–1703 (2024).

30. A. Schwab, R. Levato, M. D’Este, S. Piluso, D. Eglin, J. Malda, Printability and shape fidelity of bioinks in 3D bioprinting. Chem. Rev. 120, 11028–11055 (2020).

31. R. N. Shamma, R. H. Sayed, H. Madry, N. S. El Sayed, M. Cucchiarini, Triblock copolymer bioinks in hydrogel three-dimensional printing for regenerative medicine: A focus on Pluronic F127. Tissue Eng. Part B Rev. 28, 451–463 (2022).

32. N. S. Rojas-Higuera, A. M. Pava-Sánchez, D. L. Pinzón Rangel, L. A. Díaz-Ariza, B. Quevedo-Hidalgo, A. M. Pedroza-Rodríguez, Bio-transformed sawdust by white rot fungi used as a carrier for plant growth-promoting bacteria. Eur. J. Wood Wood Prod. 75, 263–273 (2017).

33. M. A. Shakir, M. I. Ahmad, Y. Yusup, M. Rafatullah, From waste to wealth: converting rubber wood sawdust into green mycelium-based composite. Biomass Convers. Biorefin., doi: 10.1007/s13399-023-05113-9 (2023).

34. A. J. Ragauskas, G. T. Beckham, M. J. Biddy, R. Chandra, F. Chen, M. F. Davis, B. H. Davison, R. A. Dixon, P. Gilna, M. Keller, P. Langan, A. K. Naskar, J. N. Saddler, T. J. Tschaplinski, G. A. Tuskan, C. E. Wyman, Lignin valorization: improving lignin processing in the biorefinery. Science 344, 1246843 (2014).

35. M. Sydor, G. Cofta, B. Doczekalska, A. Bonenberg, Fungi in mycelium-Based Composites: Usage and recommendations. Materials (Basel) 15, 6283 (2022).

36. W. Aiduang, J. Kumla, S. Srinuanpan, W. Thamjaree, S. Lumyong, N. Suwannarach, Mechanical, physical, and chemical properties of mycelium-based composites produced from various lignocellulosic residues and fungal species. J. Fungi (Basel) 8, 1125 (2022).

37. N. Attias, O. Danai, T. Abitbol, E. Tarazi, N. Ezov, I. Pereman, Y. J. Grobman, Mycelium bio-composites in industrial design and architecture: Comparative review and experimental analysis. J. Clean. Prod. 246, 119037 (2020).

38. N. Lin, A. Taghizadehmakoei, L. Polovina, I. McLean, J. C. Santana-Martínez, C. Naese, C. Moraes, S. J. Hallam, J. Dahmen, 3D bioprinting of food grade hydrogel infused with living pleurotus ostreatus mycelium in non-sterile conditions. ACS Appl. Bio Mater. 7, 2982–2992 (2024).

39. N. A. Gow, G. M. Gadd, *Growing Fungus* (Springer Science & Business Media, New York, NY, 2007).

40. K. Aleklett, P. Ohlsson, M. Bengtsson, E. C. Hammer, Fungal foraging behaviour and hyphal space exploration in micro-structured Soil Chips. ISME J. 15, 1782–1793 (2021).

41. J. R. M. Tuyishime, E. C. Hammer, M. Pla-Ferriol, K. Thånell, C. Alwmark, S. van Velzen, D. Floudas, R. Platakyte, M. Obst, H. Zou, Nanoscale characterization of fungal-induced CaCO3 precipitation: Implications for self-healing concrete. ACS Appl. Mater. Interfaces 17, 37648–37656 (2025).

42. S. Bindschedler, G. Cailleau, É. Verrecchia, Role of fungi in the biomineralization of calcite. Minerals 6, 41 (2016).

43. A. Van Wylick, A. V. Monclaro, E. Elsacker, S. Vandelook, H. Rahier, L. De Laet, D. Cannella, E. Peeters, A review on the potential of filamentous fungi for microbial self-healing of concrete. Fungal Biol. Biotechnol. 8, 16 (2021).

44. F. Natalio, R. Maria, Microbial biomineralization of alkaline earth metal carbonates on 3D-printed surfaces. ACS Appl. Mater. Interfaces 16, 6327–6336 (2024).

45. S. B. Mukkamala, C. E. Anson, A. K. Powell, Modelling calcium carbonate biomineralisation processes. J. Inorg. Biochem. 100, 1128–1138 (2006).

46. I. M. Van Meerbeek, J. M. Lenhardt, W. Small, T. M. Bryson, E. B. Duoss, T. H. Weisgraber, Compressive properties of silicone Bouligand structures. MRS Bull. 48, 325–331 (2023).

47. J.-H. Zhao, S.-F. Ma, X.-Y. Wei, Finite Element Analysis of Femur Stress under Bending Moment and Compression Load (IEEE, 2009)2009 2nd International Conference on Biomedical Engineering and Informatics.

48. E. Elsacker, M. Zhang, M. DadeLRobertson, Fungal engineered living materials: The viability of pure mycelium materials with selfLhealing functionalities. Adv. Funct. Mater. 33, 2301875 (2023).

49. Y. Cai, H. Yuan, C. Ronders, V. Mottini, K. Singh, L. Xing, I. Singh, D. Fu, K. Zhao, L. Heller, K. Nguyen, B. Waller, T. Wang, G. Bonito, J. Li, Mycoelectronics: Bioprinted living fungal bioelectronics for artificial sensation. Proc. Natl. Acad. Sci. U. S. A. 123, e2530456123 (2026).

50. S. Heid, A. R. Boccaccini, Advancing bioinks for 3D bioprinting using reactive fillers: A review. Acta Biomater. 113, 1–22 (2020).

51. L. Xiaorui, Z. Fuyin, W. Xudong, G. Xuezheng, Z. Shudong, L. Hui, D. Dandan, L. Yubing, W. Lizhen, F. Yubo, 1Biomaterial inks for extrusion-based 3D bioprinting: Property, classification, modification, and selection. Int. J. Bioprinting 9, 649 (2023).

