## Supporting Information for "Biosynthetic additive manufacturing through mycelium-bacteria consortia"

Hang Yuan *et al.*

#### **This PDF file includes:**

Materials and Methods

Supplementary Text

Figs. S1 to S23

Tables S1 to S5

References (1 to 15)

#### **Other Supplementary Materials for this manuscript include the following:**

Movies S1 to S8

### MATERIALS AND METHODS

#### Materials

Soybean peptone (CAS 73049-73-7), Pluronic F127 (F127, CAS 9003-11-6), tryptone (CAS 91079-40-2), yeast extract (CAS 8013-01-2), calcofluor white (CFW, CAS 18909-50-5), urea ($\text{CO}(\text{NH}_2)_2$, CAS 57-13-6), sodium chloride ($\text{NaCl}$, CAS 7647-14-5), ammonium chloride ($\text{NH}_4\text{Cl}$, CAS 12125-02-9), sodium phosphate dibasic ($\text{Na}_2\text{HPO}_4$, CAS 7558-79-4), formaldehyde ($\text{CH}_2\text{O}$, 37 wt%, CAS 50-00-0), and phosphate-buffered saline (PBS, pH 7.2) were purchased from Sigma-Aldrich, USA. Malt extract (CAS 8002-48-0) and κ-carrageenan (CAS 11114-20-8) were purchased from Research Products International, USA. Agar (CAS 9002-18-0) and calcium chloride dihydrate ($\text{CaCl}_2 \cdot 2\text{H}_2\text{O}$, CAS 10035-04-8) were purchased from Carolina Biological Supply Co., USA. Sodium hydroxide ($\text{NaOH}$, 98.0%, CAS 1310-73-2), hydrochloric acid ($\text{HCl}$, 35 wt%, CAS 7647-01-0), and Alizarin Red S sodium salt (ARS, 1% w/v, CAS 130-22-3) were purchased from Thermo Fisher Scientific, USA. Microcrystalline cellulose (MCC, CAS 9004-34-6, Loudwolf Industrial & Scientific, USA), carboxymethyl cellulose (CMC, CAS 9000-11-7, La Tiendita Essentials, USA), and propidium iodide (PI, CAS 25535-16-4, Life Technologies, USA) were used as received. Glutaraldehyde (25% aqueous solution, CAS 111-30-8), paraformaldehyde (16% aqueous solution, CAS 30525-89-4), osmium tetroxide (CAS 20816-12-0), uranyl acetate (CAS 6159-44-0), acetone (CAS 67-64-1) and Spurr resin (CAS 52368-54-4) were purchased from Electron Microscopy Sciences, USA. *Trametes versicolor* (*T. versicolor*) sawdust spawn and *Ganoderma sessile* (*G. sessile*) were obtained from Field & Forest Products, USA. *Sporosarcina pasteurii* (*S. pasteurii*, ATCC #11859) was obtained from the American Type Culture Collection, USA. Deionized (DI) water used for all solution preparation and sample processing was obtained from a Thermo Scientific water purification system (Thermo Fisher Scientific Inc., USA).

#### Preparation of mycelium inks

To prepare the F127-based mycelium ink, 30% (w/v) F127 powder was dissolved in pre-chilled (4 °C) 2% (w/v) malt extract liquid medium supplemented with 0.5% (w/v) peptone (pH = 6.8) by gentle stirring in an ice bath until fully transparent. The solution was sterilized by filtration through a 0.22 $\mu\text{m}$ polyethersulfone (PES) syringe filter (Millex-GP, SLGP033RS, MilliporeSigma, USA) under aseptic conditions. Separately, *G. sessile* was cultured in 50 mL of the same malt extract liquid medium for 5 days. After centrifugation at 4000 rpm for 10 minutes, the supernatant was discarded and the collected mycelium was gently mixed into 50 mL of the sterile F127 solution to achieve a uniform suspension. The resulting ink was stored at 4 °C to maintain its liquid state and was gently stirred before use to prevent mycelial sedimentation and ensure uniform dispersion.

Malt extract agar (MEA)-based mycelium ink was prepared by modifying standard MEA medium with rheology modifiers, as adapted from a previous study (1). A solution was prepared by dispersing 6% (w/v) malt extract, 4.5% (w/v) agar, 1.5% (w/v) κ-carrageenan, and 0.5% (w/v) peptone in DI water. Separately, 1% (w/v) MCC and 1% (w/v) CMC were pre-dispersed using a DAC 1100 mixer (FlackTek Inc., USA) at 1600 rpm for 4 minutes. The MCC/CMC suspension was then combined with the remaining components and homogenized using an immersion blender (Ganiza Inc., USA). The resulting mixture was sterilized by autoclaving at

121 °C for 20 minutes. After cooling and solidification, the MEA-based pure ink was mechanically disrupted into a paste-like consistency using the blender and spread evenly into Petri dishes. A $1 \times 1 \text{ cm}^2$ agar plug of *G. sessile* was placed on the surface, and the plates were incubated at 28 °C, under dark conditions and 95% relative humidity (RH), for 7–10 days until the surface was fully colonized by mycelium. The dense surface layer and plug were removed, and the remaining ink was re-homogenized using the mixer at 1600 rpm for 2 minutes before further use.

The sawdust-based mycelium ink was prepared using a formulation similar to the MEA-based ink, excluding malt extract and peptone. All other component concentrations and preparation procedures remained consistent. Before use, the surface layer of dense mycelial growth on the sawdust was removed to prevent nozzle clogging during printing. The autoclaved and solidified medium was fragmented and combined with 25% (w/w) *T. versicolor* sawdust spawn. The mixture was then processed using the DAC 1100 mixer at 1600 rpm for 2 minutes, with 15 mm grinding media added to assist homogenization. As a control, sawdust containing *T. versicolor* mycelium was autoclaved before mixing, while all other steps were kept identical.

##### Rheology of mycelium inks

The rheological behaviour of four ink formulations was measured, including F127-based mycelium ink, MEA-based pure ink, MEA-based mycelium ink, and sawdust-based mycelium ink. All measurements were conducted using a Discovery HR-2 rheometer (TA Instruments, USA) equipped with a 40 mm parallel plate geometry and a 1000 $\mu\text{m}$ gap. All measurements were performed at 25 °C, and data were processed using TRIOS software (TA Instruments, USA). Apparent viscosity was recorded over a shear rate range of 0.1 to 100 $\text{s}^{-1}$ to assess shear-thinning behaviour under steady shear. Oscillatory strain sweep tests were performed at a constant angular frequency of 10 rad/s with strain amplitudes ranging from 0.1% to 100% to determine the storage modulus ($G'$), the loss modulus ($G''$), and the yield point corresponding to the crossover of $G'$ and $G''$. Shear recovery behaviour was assessed by alternating oscillatory (1% strain, 10 rad/s) and steady shear at 1 $\text{s}^{-1}$ in repeated cycles. Each cycle included 60 seconds of oscillatory measurement to establish the baseline moduli, followed by 30 seconds of steady shear to simulate extrusion, and concluded with a return to oscillatory mode to monitor recovery.

##### Preparation of growth media and fungal culture

Commercially obtained *G. sessile* was inoculated on MEA medium for initial culture and routine subculturing. To prepare the MEA medium, 2.0% (w/v) malt extract, 0.5% (w/v) peptone, and 1.5% (w/v) agar were dissolved in deionized (DI) water. The pH of the solution was adjusted to 6.8 using 1 M NaOH and 1 M HCl. The mixture was transferred into borosilicate glass bottles, loosely capped, and the bottle openings were covered with aluminum foil to prevent contamination during autoclaving. Sterilization was performed at 121 °C for 20 minutes using a high-pressure steam autoclave (Consolidated Sterilizer Systems, USA). After cooling, the sterilized medium was poured into sterile Petri dishes and allowed to solidify.

For subculturing, a $1 \times 1 \text{ cm}^2$ agar plug fully covered with mycelium was excised and transferred to the center of a 2% MEA plate (2% malt extract) every four weeks. The inoculated plates were incubated at 28 °C, under dark conditions, with 95% RH, in an incubator (HERAcell VIOS 160i, Thermo Fisher Scientific Inc., Waltham, MA, USA).

Malt extract liquid medium was prepared by dissolving 2.0% (w/v) malt extract and 0.5% (w/v) peptone in DI water (pH = 6.8). All sterilization procedures were consistent with those described for the MEA medium. Liquid cultures were initiated by transferring 1 × 1 cm<sup>2</sup> agar plugs fully colonized with *G. sessile* mycelium into 50 mL of sterile liquid medium in 250 mL Erlenmeyer flasks. The incubation conditions for liquid cultures were identical to those used for MEA plates (28 °C, ~95% RH, in the dark).

*T. versicolor* was used in the form of commercially prepared sawdust spawn. According to the manufacturer's specifications, the hardwood sawdust was sterilized at 121 °C for 30 minutes, cooled, and then sealed in 0.5 µm filter patch bags with gussets to allow gas exchange while preventing microbial contamination. The mycelium was introduced into the substrate using mechanical inoculation tools, followed by incubation to achieve full colonization. The sawdust spawn was stored at 4 °C until use.

##### Mycelial growth behaviour and bridging across air gaps

To investigate the influence of malt extract concentration on mycelial growth behaviour, MEA plates were prepared with malt extract concentrations ranging from 0% to 10% (w/v). A 1 × 1 cm<sup>2</sup> *G. sessile* mycelium-covered agar plug, previously cultured on a 2% MEA plate, was placed at the center of each plate. The plates were then incubated at 28 °C under dark conditions with 95% RH. After five days of growth, plates at certain concentrations exhibited nearly complete surface colonization. Mycelial expansion was assessed by measuring both radial growth and pixel intensity. Radial growth was defined as the distance from the center of the inoculum to the edge of the mycelial colony. Pixel intensity was analyzed by randomly selecting a 300 px × 300 px region (approximately 8.8 mm × 8.8 mm) within the colonized area. Intensity values ranged from 0 (black) to 255 (white), with denser mycelial regions yielding higher values. For each plate, five measurements were performed and averaged, and five replicate plates were analyzed for each concentration. All measurements were conducted using ImageJ software (National Institutes of Health, Bethesda, MD, USA). All images were captured inside a controlled LED photo studio (LB12, Takerers, USA) using a Canon EOS R8 camera (Canon Inc., Tokyo, Japan) equipped with a macro-objective (RF35mm F1.8 Macro STM, Canon).

The bridging capability of mycelium, defined as the ability of hyphae to grow laterally across unsupported air gaps between adjacent printed filaments, was evaluated for *T. versicolor* and *G. sessile*. For *T. versicolor*, sawdust-based mycelium ink was extruded into filaments approximately 6 mm in diameter using a 3D PotterBot 10 PRO printer (3D Potter, USA), forming parallel lines separated by air gaps ranging from 1 mm to 7 mm, in 1 mm increments. For *G. sessile*, MEA-based mycelium ink was printed using a Brinter ONE bioprinter (Brinter Ltd., Finland) fitted with a 1.6-mm nozzle to generate air gaps of 0.5 mm to 2.5 mm, in 0.5 mm increments. All samples were incubated in the dark under conditions of 28 °C and 95% RH for 7 days (*T. versicolor*) or 4 days (*G. sessile*). Bridging was considered successful when the mycelium visibly spanned and connected the majority of the gap area between adjacent filaments. Hyphal growth across the gaps was monitored daily to assess bridging performance.

#### 3D printing of mycelium-based mechanical specimens

Mechanical test specimens were fabricated using a 3D PotterBot 10 PRO printer. The 3D structures were designed in SolidWorks 2024 (Dassault Systèmes, France), and exported as stereolithography (STL) files. Printing paths were generated as geometric (G)-code using UltiMaker Cura slicing software (version 5.9.1, UltiMaker B.V., Netherlands). All printing was conducted at room temperature with a constant printing speed of 20 mm/s. All constructs were printed onto sterilized polyvinyl chloride (PVC) sheets to prevent contamination. Compression test specimens were printed with a square footprint and stacked to form 16-layer structures. A single-shell outer wall was printed to enclose the internal filament pattern. Inside the shell, sawdust-based mycelium ink was extruded as filaments using a 6-mm anodized aluminum nozzle (3D Potter, USA). Filaments in adjacent layers were aligned in parallel and rotated at specific spiral angles (15°, 30°, 45°, and 90°), forming controlled air gaps (1, 2 and 3 mm). In contrast, three-point bending test specimens consisted of 4 layers, with all filaments oriented at 90° and spaced 1 mm apart. After printing, all samples were incubated at 28 °C, in the dark, at 95% RH for 7 days, and subsequently dried to a constant weight in a convection oven (Precision™, Thermo Scientific, USA) at 70 °C.

#### Mechanical compression and three-point bending tests

Uniaxial compression tests were conducted using a universal testing machine (Model SFM-20, United Calibration Corp., Huntington Beach, CA, USA) equipped with a 20,000-lb load cell under 25 °C conditions. Mycelium-based specimens were prepared and tested in groups of five per condition. For each sample, the length, width, and height were measured at five positions using a digital caliper, and the average values were used to calculate the cross-sectional area and original height. A constant crosshead speed of 6 mm/min was maintained until structural failure occurred. Compressive strain ($\varepsilon$) was calculated as the ratio of displacement ($\Delta L$) to the initial height ($L_0$), and compressive stress ($\sigma$) was obtained by dividing the applied force ($F$) by the specimen's cross-sectional area ($A$), as shown below:

$$\varepsilon = \frac{\Delta L}{L_0} \quad (S1)$$

$$\sigma = \frac{F}{A} \quad (S2)$$

The Young's modulus ($E$) was obtained by the linear fitting of the stress-strain curve in the 0–1% strain region, and the ultimate compressive strength ($\sigma_U$) was determined as the maximum stress before failure, as shown below:

$$E = \left. \frac{d\sigma}{d\varepsilon} \right|_{\varepsilon=0-0.01} \quad (S3)$$

$$\sigma_U = \max(\sigma) \quad (S4)$$

The bulk density ($\rho$) of each dried sample was calculated by dividing its dry mass ($m$) by its geometric volume ($V$) using the following equation:

$$\rho = \frac{m}{V} \quad (\text{S5})$$

$$V = L_0 \times A \quad (\text{S6})$$

Three-point bending tests were performed on rectangular specimens using the same universal testing machine and software as described above, equipped with a 100-lb load cell. The support span was fixed at 57 mm, and the loading rate was set to 6 mm/min. All measurements were conducted at 25 °C. The flexural stress ($\sigma$) and strain ($\varepsilon$) were calculated using the following equations:

$$\sigma = \frac{3FL}{2bd^2} \quad (\text{S7})$$

$$\varepsilon = \frac{6Dd}{L^2} \quad (\text{S8})$$

where $F$ is the applied force, $L$ is the support span, $b$ and $d$ are the width and thickness of the specimen, and $D$ is the midspan deflection.

The flexural modulus ($E_f$) was obtained by fitting the initial linear portion (0–0.5% strain) of the load–displacement curve:

$$E_f = \frac{L^3 m}{4bd^3} \quad (\text{S9})$$

where $m$ is the slope of the load-displacement curve in the linear region. The flexural strength ($\sigma_U$) was defined as the maximum stress before failure.

All data analyses and curve fittings were performed using Origin 2021 (OriginLab Corporation, Northampton, MA, USA). To obtain interpolated average curves, the strain range was first standardized across all samples based on their common overlap, and interpolation was conducted at a fixed interval of 0.02% strain. Curves were then averaged point-by-point to generate representative trends for each group.

##### Bacteria culturing and biomineralization of mycelium scaffolds

*S. pasteurii* was cultured in an alkaline peptone yeast extract (APY) medium composed of 1.0% (w/v) tryptone, 0.5% (w/v) yeast extract, 0.5% (w/v) NaCl, 0.5% (w/v) NH<sub>4</sub>Cl, and 0.1% (w/v) Na<sub>2</sub>HPO<sub>4</sub>. The pH was adjusted to 8.0 using 1 M NaOH and 1 M HCl before autoclaving at 121 °C for 20 minutes. All media and reagents were prepared using DI water. Frozen primary stocks of *S. pasteurii* were thawed and transferred into 50 mL of sterile APY medium, followed by incubation at 28 °C and 100 rpm in an orbital shaking incubator (MaxQ 6000, Thermo Fisher Scientific, USA). To monitor bacterial growth, 100 μL of the culture was sampled daily and diluted in fresh APY medium. The optical density (OD) at 600 nm was then measured using a NanoDrop OneC spectrophotometer (Thermo Fisher Scientific, USA) with 1.5 mL disposable semi-micro cuvettes (BRAND, Germany). Each measurement was performed five times and the average value was recorded.

For co-culturing with mycelium scaffolds, *S. pasteurii* cultures were harvested at the late exponential phase ($OD_{600} \approx 0.6\text{--}0.8$), centrifuged at 4000 rpm for 10 minutes using an LD-3 Electronic Centrifuge (China), and resuspended in a fresh APY medium. Mycelium scaffolds, which had been pre-incubated at 28 °C and 95% RH for 7 days to allow full surface colonization, were transferred into sterilized glass beakers and completely submerged in a large excess volume of the bacterial suspension (maintaining a liquid-to-solid volume ratio of $> 2:1$). The beakers were loosely covered with aluminum foil and incubated at 28 °C while shaking at 100 rpm for 24 hours.

To induce biomineralization, a 1:1 (v/v) replacement of the bacterial suspension with calcium-based mineralization medium (CBMM) was performed. The CBMM was composed of 2.0% (w/v) urea, 1.0% (w/v) $\text{CaCl}_2 \cdot 2\text{H}_2\text{O}$, 0.1% (w/v) yeast extract, and 0.1% (w/v) $\text{Na}_2\text{HPO}_4$, with the pH adjusted to 8.0 using 1 M NaOH. The CBMM was adapted from previously reported biomineralization protocols (2, 3). Due to the thermal instability of urea, all components except urea were dissolved in DI water and sterilized by autoclaving at 121 °C for 20 minutes. Urea was dissolved separately and sterilized by filtering through a 0.22 $\mu\text{m}$ PES syringe filter under aseptic conditions, and then mixed with the autoclaved solution before use. The scaffolds were incubated for an additional 48 hours under the same conditions. Finally, the supernatant and any surrounding precipitates were carefully removed, and the scaffolds were dried in a convection oven at 70 °C until constant weight.

##### Preparation of self-healing and self-assembly mycelial scaffolds

Self-healing and self-assembly mycelial scaffolds were prepared using the same printing, drying, biomineralization, and compression conditions as other compression specimens ($\alpha = 90^\circ$, $g = 1\text{ mm}$). For the self-healing samples, after printing, an autoclaved stainless-steel scalpel was used to make a single scratch on one surface, with a length of approximately five filament layers. The scratched samples were then incubated and dried for testing. For the self-assembly samples, each scaffold was printed as two equal-height parts, which were incubated separately for 7 days and then manually placed together and further incubated for another 7 days to allow them to self-assemble into a single integrated structure before drying.

##### 3D printing of mycelium inks and self-assembly of printed structures

The mycelium inks were printed using a Brinter ONE bioprinter equipped with a pneumatic extrusion head (Pneuma Tool Pro, Brinter Ltd., Finland). F127-based and MEA-based mycelium inks were printed through 18-gauge (0.84 mm) and 14-gauge (1.6 mm) precision tips (Nordson EFD, USA), respectively. Printing was performed at a speed of 6 mm/s directly onto sterile glass substrates (Fisherbrand, Fisher Scientific, USA). To ensure sterility, the bioprinter chamber features an internal UV-C system for print bed decontamination before printing. The stored F127-based ink was gently loaded from 4 °C into sterile cartridges (Brinter Ltd., Finland) by slowly dispensing along the inner wall to minimize air bubble formation. The filled cartridges were then left undisturbed at room temperature for 5 minutes to allow the ink to be printable.

Unless otherwise stated, the printing conditions for the sawdust-based mycelium ink were consistent with those used for fabricating mechanical testing specimens. To support the fabrication of overhanging structures using freshly extruded sawdust-based inks, temporary structural support was required to prevent sagging or collapse before the structure was stabilized through mycelial growth and drying. As shown in the accompanying image, scaffolds were fabricated using a desktop FDM 3D printer (Ender-3, Creality, China) with polylactic acid (PLA) filament (Elegoo, China), and used as removable supports. The PLA frames were detached from the dried mycelium structures.

Each printed mycelium component was initially incubated for 7 days at 28 °C and 95% RH, under dark conditions, to allow for sufficient surface colonization. After incubation, the components were either manually positioned adjacent to one another or temporarily secured in a predefined arrangement using external supports. The assembled structures were then incubated under the same conditions for an additional 7 days. During this second incubation period, the individual mycelium structures exhibited spontaneous self-assembly behaviour through hyphal extension into neighbouring units. This process was primarily driven by the directional growth of hyphae from the edge of one structure toward adjacent ones, resulting in physical interconnection across small air gaps (1). The newly formed mycelial network effectively bridged discrete printed units, forming a continuous and integrated construct without the use of external adhesives.

##### Confocal laser scanning microscopy of co-cultured mycelium and bacteria

Fungal samples of *T. versicolor* or *G. sessile* were cut into 5-10 mm fragments using sterile tools. *S. pasteurii* cultures were harvested at the late exponential phase ($OD_{600} \approx 0.6-0.8$), centrifuged at 4000 rpm for 10 min using a centrifuge, and resuspended in fresh APY media for co-culturing. The fungal fragments were immersed in the bacterial suspension in sterile conical flasks sealed with aluminum foil and incubated at 28°C while shaking at 100 rpm for 24 hours using an orbital shaking incubator. After co-culturing, samples were fixed in 4% (w/v) formaldehyde diluted in 0.1M PBS for 20 min at room temperature, followed by two washes with PBS to remove residual fixatives. Fixed samples were stained with 0.25 g/L CFW for fungal hyphae and 0.1 g/L PI for bacterial cells. Staining was performed in PBS at room temperature for 15 min in the dark, followed by two PBS washes.

To visualize mineralization, mycelium samples co-cultured with *S. pasteurii* were incubated in CBMM to induce *in situ* precipitation of calcium carbonate. Fixed samples were then stained with 0.25 g/L CFW and 1% (w/v) ARS to label mineralized calcium carbonate. Staining was performed at room temperature in the dark for 15 min without subsequent PBS washes to preserve the hydrated mineral phase. No bacterial staining was applied under these conditions to avoid spectral overlap during imaging.

Samples were mounted in PBS on standard 75 mm × 25 mm microscope slides and covered with #1.5 (0.17 mm) coverslips (Menzel-Gläser, Thermo Fisher Scientific) for imaging. Confocal laser scanning microscopy (CLSM) imaging was performed on an upright microscope (Eclipse Ni-U) equipped with a laser scanning system (C2+, Nikon Instruments Inc., Japan) and a 20×/0.75 NA objective (Plan Apo λ). CFW, PI, and ARS were excited at 405 nm, 561 nm, and 561 nm, respectively, and detected using 445/35 nm and 600/50 nm bandpass filters. Fluorescence signals were pseudo-colored using look-up tables: CFW in cyan, PI in red, and ARS in gray. Images were acquired with 2× digital zoom and 0.16 μm pixel size. Z-stack images were acquired using NIS-Elements software (version 5.42.03, Nikon Instruments Inc., Japan) with a step size of 1.00 μm, and rendered into 3D projections. The total acquisition depth varied slightly between samples and was adjusted accordingly based on actual imaging. Channels were captured sequentially with no transmitted light channel. Only linear brightness and contrast adjustments were uniformly applied across each image using NIS-Elements.

##### Scanning electron microscopy and energy dispersive X-ray microanalysis

Dried samples used for mechanical testing, including both unmineralized and mineralized mycelium scaffolds, were imaged using a JEOL JSM-6610LV scanning electron microscope (SEM) equipped with a tungsten hairpin emitter (JEOL Ltd., Tokyo, Japan). Calcium carbonate layers were collected from the surface of dried mineralized mycelium scaffolds and imaged using the same procedure. An accelerating voltage of 12 kV was applied, and images were acquired using a secondary electron (SE1) detector. Before imaging, samples were mounted on aluminum stubs using high-vacuum carbon tape (SPI Supplies, West Chester, PA, USA). Cross-sectional samples were prepared using a low-speed diamond saw (Model 650, South Bay Technology Inc., San Clemente, CA, USA). All samples were coated with a thin (~10 nm) layer of osmium using a Tennant20 osmium chemical vapor deposition coater (Meiwafosis Co., Ltd., Osaka, Japan) to enhance conductivity and image quality.

Elemental analysis and mapping were performed using energy dispersive X-ray spectroscopy (EDS) with an Oxford Instruments AZtec system (software version 3.1; Oxford Instruments, High Wycombe, Bucks, UK) integrated with the JEOL 6610LV SEM. A 20 mm<sup>2</sup> silicon drift detector with an ultra-thin window was used to collect EDS signals.

##### Powder X-ray diffraction and Fourier-transform infrared spectroscopy

Dried unmineralized and mineralized mycelium scaffolds were ground into fine powders using an agate mortar and pestle to ensure homogeneity. Powdered samples were loaded onto a poly(methyl methacrylate) specimen holder. Powder X-ray diffraction (PXRD) patterns were collected at room temperature using a Cu-Kα<sub>1</sub> radiation source (emission wavelength $\lambda = 1.54060 \text{ \AA}$) (Bruker AXS GmbH, Karlsruhe, Germany). A LynxEye 1D detector operating in continuous position-sensitive detector fast mode was used for data acquisition. The X-ray generator was operated at 40 kV and 40 mA. Diffraction data were recorded over a 2θ range of 15° to 60°, with a step size of 0.02° and a counting time of 0.05 s per step. Phase identification and peak assignment were carried out using JADE 9 software (Materials Data Inc., Livermore, CA, USA).

A separate portion of the same dried mycelium scaffold powders used for PXRD was utilized for Fourier-transform infrared spectroscopy (FT-IR) analysis to ensure consistent sample origin. Spectra were collected using a Spectrum Two FT-IR spectrometer equipped with an attenuated total reflectance (ATR) accessory (PerkinElmer Inc., Waltham, MA, USA). Samples were placed directly onto the ATR crystal with gentle pressure. Data were recorded over the range of 4000-500 $\text{cm}^{-1}$ with a spectral resolution of 4 $\text{cm}^{-1}$, averaging 32 scans per sample.

##### Micro-computed tomography

Micro-computed tomography (MicroCT) was conducted using the Quantum GX system (PerkinElmer, USA) to non-destructively visualize the internal structures of mineralized and unmineralized *T. versicolor* mycelium scaffolds. Samples were scanned in a dry state using the standard mode, with an X-ray tube voltage of 90 kV, current of 88 $\mu\text{A}$, and a field of view (FOV) of 72 mm × 72 mm. The voxel size was set to 144 $\mu\text{m}$, and each scan lasted 18 seconds. Reconstruction was performed at the same resolution using the system's integrated software. Reconstructed cross-sectional images were displayed using the original contrast mode. To enable consistent grayscale interpretation, a uniform window level setting (center = -350 a.u., width = 900 a.u.) was applied to both samples during image rendering. This approach ensured consistent grayscale interpretation across conditions. Representative slices were extracted to illustrate the distribution of mycelial networks and mineral deposits. After 8-bit conversion, the mpl-inferno LUT was applied in Fiji (National Institutes of Health, USA) for pseudo-color rendering.

##### Transmission electron microscopy

Samples were initially fixed in a solution containing 2.5% (w/v) glutaraldehyde and 2.0% paraformaldehyde (w/v) prepared in 0.1 M PBS. After initial fixation, the samples were rinsed with a 0.1 M phosphate buffer and post-fixed with 1% (w/v) osmium tetroxide in a 0.1 M phosphate buffer. Samples were then block-stained with 1% (w/v) uranyl acetate, dehydrated through a graded acetone series, and subsequently infiltrated and embedded in Spurr resin. Ultrathin sections (70 nm) were prepared using an EM UC7 ultramicrotome (Leica Microsystems, Germany), and post-stained with uranyl acetate and lead citrate. Transmission electron microscopy (TEM) images were acquired using a JEOL 1400Flash microscope (JEOL Ltd., Japan) operated at an accelerating voltage of 100 kV.

##### Compression simulation of the Bouligand structure

In this study, compression simulations were performed on sawdust-based mycelium composite materials featuring a Bouligand structure. Three-dimensional models of the composite structure were constructed in SolidWorks 2024 (Dassault Systemes, France) and imported into ANSYS Workbench 2024 R2 (ANSYS Inc., USA) for finite element analysis (FEA). Each filament in the printed structure was modeled as a solid cylinder with a diameter corresponding to the actual nozzle diameter used during extrusion-based 3D printing. A uniform line spacing (g) of 1 mm was applied. To replicate the interlayer fusion typically observed in printed soft composites, a consistent overlapping length was incorporated between adjacent layers (4). The contact condition between overlapping regions was defined as bonded in ANSYS to simulate interfacial integrity. A small uniaxial strain of $\varepsilon = 0.0015$ was applied to ensure elastic deformation conditions. Compression simulations were conducted on Bouligand structures with interlayer deflection angles of 15°, 30°, 45°, and 90°, respectively. The effective

Young's modulus for each structure was calculated based on the simulated stress–strain data, using the same formula applied in mechanical testing.

#### Compression simulation of microscale unmineralized and mineralized mycelial networks

To further investigate the mechanical characteristics of unmineralized and mineralized mycelial networks at the microscale, we reconstructed 3D fungal structures from CLSM z-stacks of *T. versicolor* using Fiji software. The resulting STL files were imported into Rhino 8 (Robert McNeel & Associates, USA) for mesh repair and volumetric modeling. Due to the complex curvature and heterogeneity of the hyphal network, we selected a representative subvolume (24 $\mu\text{m} \times 24 \mu\text{m} \times 15 \mu\text{m}$) from the larger structure to minimize computational complexity. The choice of subvolume was based on regions with typical hyphal density and connectivity. Two comparative models were constructed using ANSYS Workbench 2024 R2. One model contained only mycelial filaments, with the pore regions treated as air. The other model represented the mineralized case, in which calcium carbonate was introduced to fill the mycelial pores using Boolean operations to simulate biomineralization. The porosity of the mineralized region was controlled to match an average value of 16.17%, which was determined by analyzing binarized 2D microCT image slices using Fiji. The average area fraction of pores across selected slices was used to estimate the porosity for the simulation. To reflect this porosity in the 3D model, spherical pores with randomized radii and spatial positions were generated and imported into SolidWorks, where Boolean subtraction was applied to remove these volumes from the mineralized region. The contact between calcium carbonate and mycelium was defined as bonded. To ensure linear elastic deformation, a small compressive displacement ($\delta = 0.001 \mu\text{m}$, corresponding to $\varepsilon \approx 0.0067\%$) was applied to the top surface, while the bottom surface was fixed (fig. S11).

The material properties used in the simulation were derived from the references (5, 6). The Young's modulus ($E$) of *T. versicolor* hyphae was approximated from the dimitic *Grifola frondosa* (maitake), based on a reported value of 561 MPa. This substitution is justified by the shared dimitic hyphal architecture and similar biochemical composition (primarily chitin-based cell walls) of both Basidiomycetes (7–9). The Poisson's ratio ($\nu$) of the mycelium was estimated to be 0.25, in line with values reported for chitin fibers (5). For calcium carbonate, the Young's modulus and Poisson's ratio were set to 33.98 GPa and 0.31, respectively, as determined for biomineralized calcite precipitated by *S. pasteurii*, and applied here to model the mineralized hyphal network (6).

Young's modulus values were measured using nanoindentation in the references and calculated using the standard reduced modulus formula (10):

$$\frac{1}{E_r} = \frac{1 - \nu_s^2}{E_s} + \frac{1 - \nu_i^2}{E_i} \quad (\text{S10})$$

where $E_r$ is the reduced modulus, $E_s$ and $\nu_s$ are the sample's Young's modulus and Poisson's ratio, and $E_i$ and $\nu_i$ are the indenter's modulus (1140 GPa) and Poisson's ratio (0.07).

#### Supplementary Text 1. Herschel–Bulkley model fitting for bioink rheology.

To quantitatively characterize the flow behaviour of the bio-inks, rheological data in the low shear rate regime ($0.1\text{--}5\text{ s}^{-1}$) were fitted to the Herschel–Bulkley model, defined as:

$$\tau = \tau_y + K \cdot \dot{\gamma}^n \quad (\text{S11})$$

where $\tau$ is the shear stress (Pa), $\tau_y$ is the yield stress (Pa), $\dot{\gamma}$ is the shear rate ($\text{s}^{-1}$), $K$ is the consistency index ($\text{Pa} \cdot \text{s}^n$), and $n$ is the flow behaviour index (dimensionless).

This model is suitable for describing non-Newtonian, yield-stress fluids with shear-thinning or shear-thickening behaviour (11, 12). The selected range ($0.1\text{--}5\text{ s}^{-1}$) reflects the low-shear conditions encountered during extrusion-based bioprinting. The good agreement between experimental data and the fitted curves supports the appropriateness of this model for evaluating ink printability and structural stability.

#### Supplementary Text 2. Representative volume element modeling with periodic boundary conditions for Bouligand structures.

To investigate the mechanical behaviour of composite structures, finite element simulations often employ representative volume elements (RVEs) with periodic boundary conditions (PBCs) to eliminate edge effects and capture intrinsic deformation responses (13). By ensuring that opposite boundaries deform identically, PBCs enable accurate evaluation of internal stress and strain distributions within repeating microstructures. Depending on the analysis objective, RVEs may contain one or multiple unit cells. In practice, deformation constraints are typically applied using fixed displacements or prescribed strains to maintain periodicity and simulate bulk-like behaviour (14).

As an example, the PBCs applied to the sawdust-based mycelium composite with a Bouligand architecture are illustrated in fig. S14A. In the case of a two-dimensional model, the structure is assumed to be periodic only along the y-axis. A prescribed displacement ($U$) is applied to the top boundary ($Yb$), while the bottom boundary ($Ya$) is fixed in the y-direction. To enforce periodicity in the x-direction, the displacements of all nodes on the lateral edges ($Xa$ and $Xb$) are coupled to be identical in the x-direction, ensuring uniform deformation across the unit cell.

The boundary conditions for the 2D case are specified as follows:

$Ya$: $UY = 0$, $UX = \text{Free}$

$Xa, Xb$: $UY = \text{Free}$, $UX$ of all nodes are coupled (equal on this edge)

$Yb$: $UX = \text{Free}$, $UY = U$ where $U = l_x \varepsilon$, with $l_x$ represents the original length of sides $Xa$ and $Xb$, $\varepsilon$ is the applied strain on the unit cell in the y-direction

For the three-dimensional model, where the structure remains periodic along the y-direction, similar PBCs are applied. The boundary conditions are defined as:

$Ya$: $UY = 0$; $UX, UZ = \text{Free}$

$Yb$: $UX, UZ = \text{Free}$, $UY = \delta$ where $\delta = l \times \varepsilon$, $\varepsilon$ is the applied strain on the unit cell in the y-direction in fig. S14B.

$Xa, Xb$: $UY, UZ$= Free; $UX$ of all nodes are coupled (equal on this edge)

$Za, Zb$: $UX, UY$= Free; $UZ$ of all nodes are coupled (equal on this edge)

These conditions ensure that the unit cell deforms in a manner consistent with the periodic microstructure of the Bouligand composite, eliminating boundary effects while enabling the extraction of effective mechanical properties under compression.

### Supplementary Figures

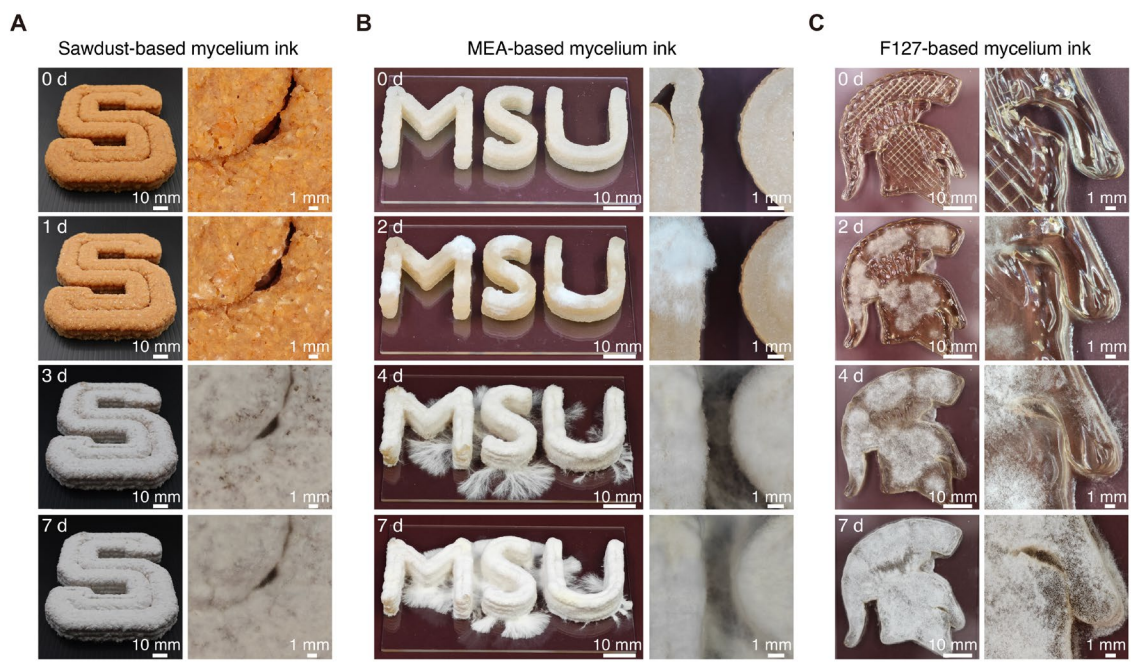

**Fig. S1.**

**Growth progression of 3D-printed mycelium scaffolds using different mycelium inks.** Representative overview (left) and zoom-in (right) images showing the growth of mycelium over time (0–7 days) in three types of printed scaffolds: **(A)** Letter “S” printed with sawdust-based mycelium ink inoculated with *T. versicolor*. **(B)** “MSU” letterforms printed with MEA-based mycelium ink inoculated with *G. sessile*. **(C)** Spartan helmet pattern printed with F127-based mycelium ink inoculated with *G. sessile*. Each row corresponds to a different cultivation day, illustrating the progression of mycelial colonization and morphological changes. Representative images from 0 d and 7 d were selected for inclusion in Fig. 2.

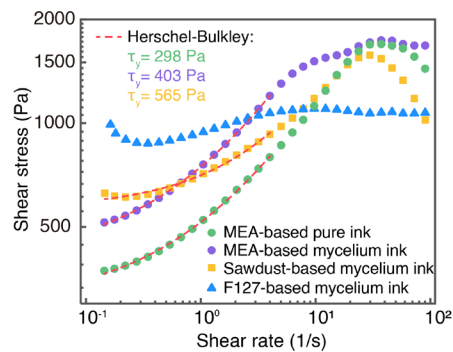

**Fig. S2.**

**Rheological behaviour of mycelium and pure inks.** Shear stress versus shear rate for mycelium inks and MEA-based pure ink without mycelium. All formulations exhibit shear-thinning behaviour. Apparent yield stress ($\tau_y$) values were obtained by fitting the data to the Herschel–Bulkley model (dashed red lines) in the low shear rate range of 0.1-5 $s^{-1}$.

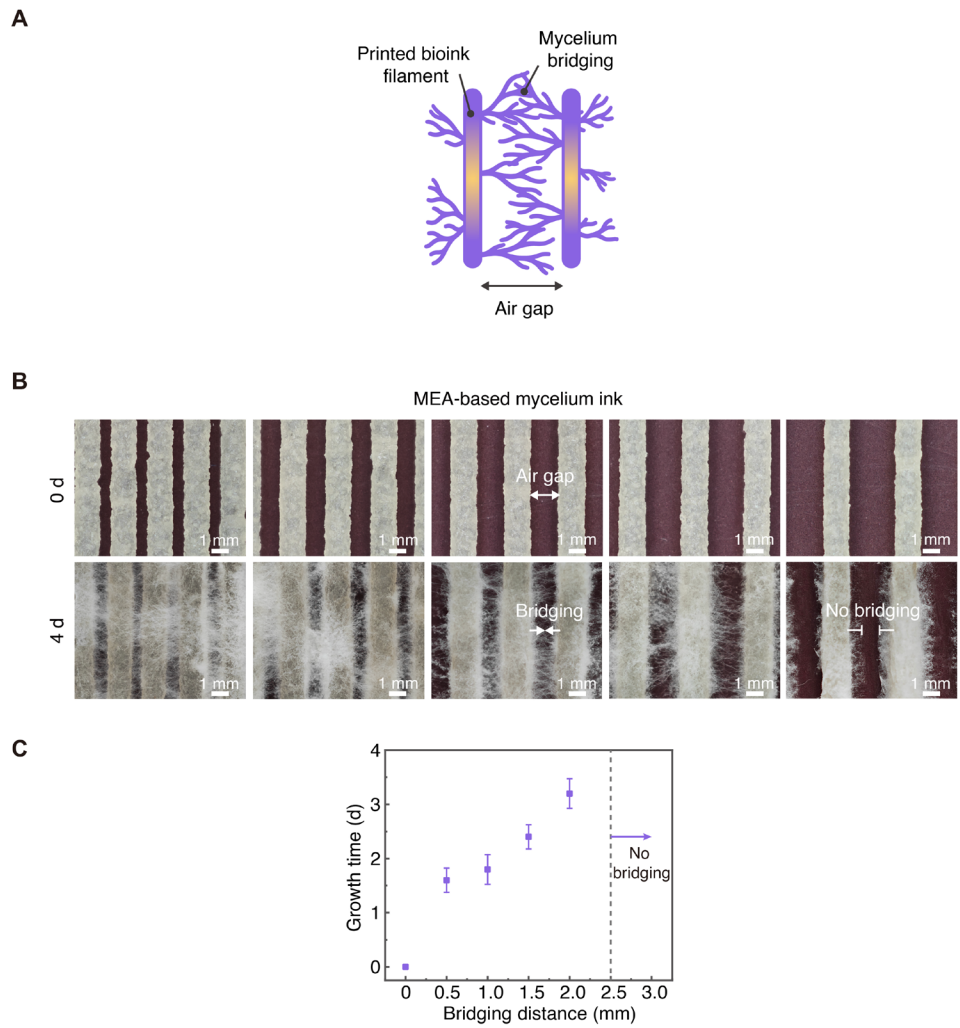

**Fig. S3.**

**Mycelial bridging across air gaps between printed filaments.** (A) Schematic illustration of mycelium growing from printed bioink filaments and bridging adjacent structures across air gaps. (B) Optical images showing the growth of *G. sessile* mycelium in MEA-based bio-ink scaffolds with varying air gap distances (0.5-2.5 mm), captured before (0 d) and after 4 days of growth (4 d). (C) Growth time required for mycelium to bridge across varying gap distances ($n = 5$). No bridging was observed when the gap exceeded 2.5 mm.

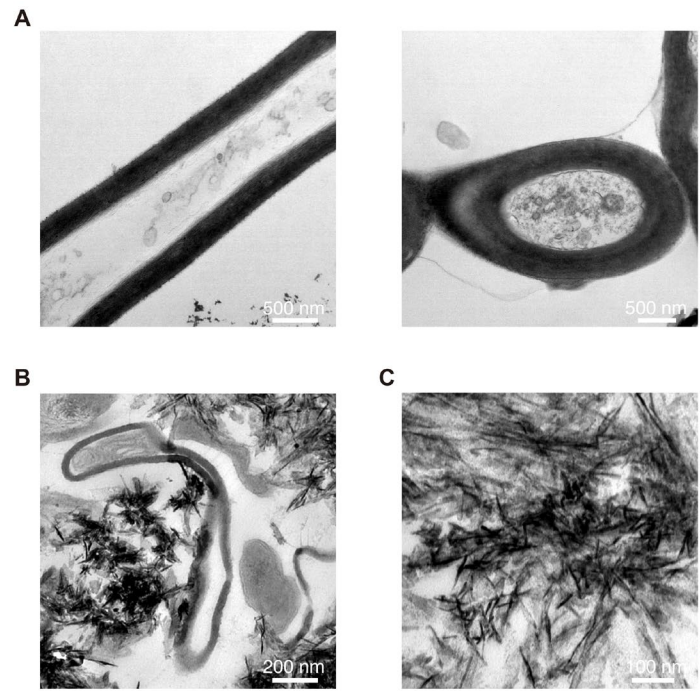

**Fig. S4.**

**TEM images of unmineralized and mineralized *T. versicolor* hyphae, and mineralization products.** (A) Left: Longitudinal section of an unmineralized hypha, showing a cell wall enclosing a lumen. Right: Transverse section of an unmineralized hypha with a lumen and wall layering. (B) TEM image of a mineralized hypha. The hypha exhibits a boundary corresponding to the cell wall, with a lighter lumen region. Mineral deposits are visible around the hyphal surface (dark crystalline regions). (C) Aggregated mineralization products.

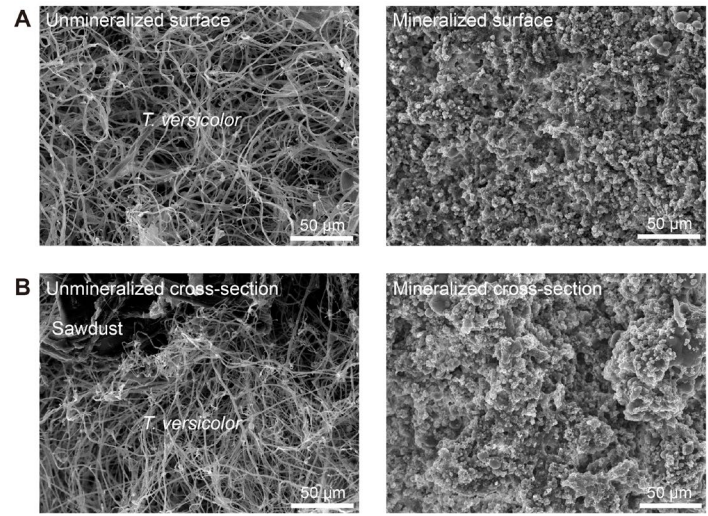

**Fig. S5.**

**SEM images showing morphological changes in *T. versicolor* mycelium scaffolds before and after biomineralization.** (A) Surface morphology of unmineralized (left) and mineralized (right) *T. versicolor* mycelium scaffolds. (B) Cross-sectional morphology of unmineralized (left) and mineralized (right) *T. versicolor* mycelium scaffolds containing sawdust. Biomineralization results in the formation of a dense, granular mineral layer that covers the originally fibrous mycelial network.

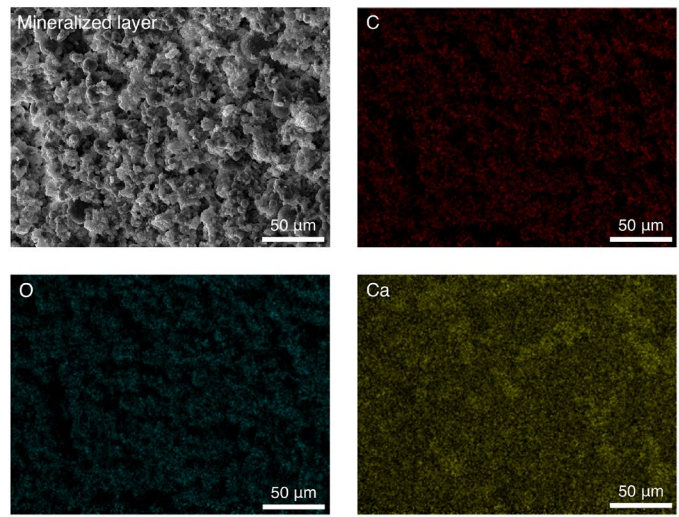

**Fig. S6.**

**Elemental mapping of the mineralized layer formed on the surface of mineralized *T. versicolor* mycelium scaffolds.** SEM image (left) and corresponding EDS elemental maps showing the distribution of carbon (C, red), oxygen (O, cyan), and calcium (Ca, yellow) in a mineralized layer peeled from the surface of a *T. versicolor* mycelium scaffold after biomineralization.

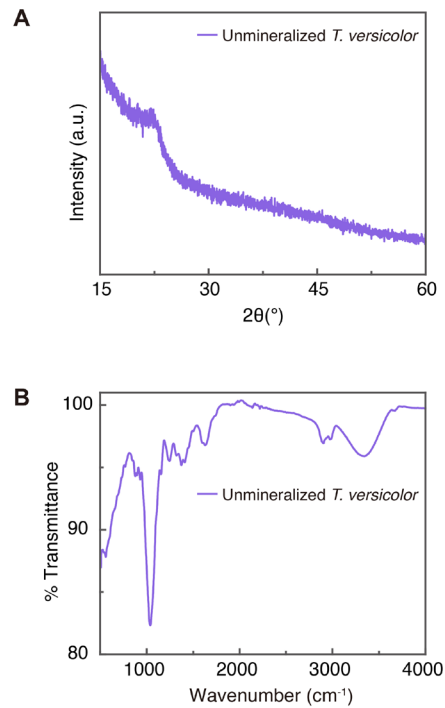

**Fig. S7.**

**PXRD and FTIR of unmineralized *T. versicolor* mycelium scaffold.** (A) PXRD pattern showing a broad, featureless curve without sharp crystalline peaks, indicating the amorphous nature of the scaffold and the absence of crystalline calcium carbonate. (B) FT-IR spectrum showing characteristic absorption bands around 1050–1100 $\text{cm}^{-1}$ (C–O–C stretching in polysaccharides), 1630–1650 $\text{cm}^{-1}$ (amide I band of proteins), and 3300–3400 $\text{cm}^{-1}$ (O–H/N–H stretching), confirming the presence of fungal cell wall components such as polysaccharides and proteins (15).

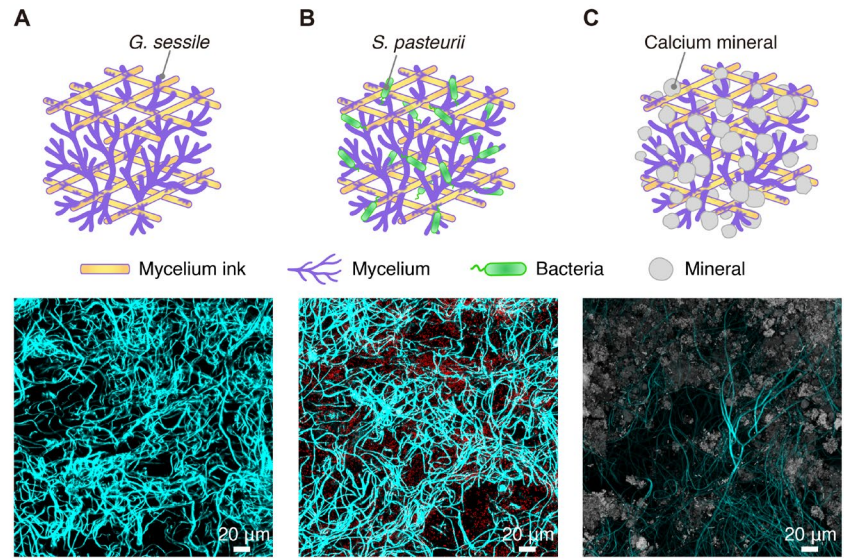

**Fig. S8.**

**CLSM images showing the mycelial scaffold before and after co-culture with bacteria and biomineralization.** (A) Mycelial network formed by *G. sessile* (shown in cyan). (B) Co-culture of *G. sessile* with *S. pasteurii*, showing fungal hyphae and membrane-compromised bacterial cells (shown in red). (C) Mycelial scaffold after calcium induction, with hydrated calcium minerals (shown in gray) precipitating along the hyphal network.

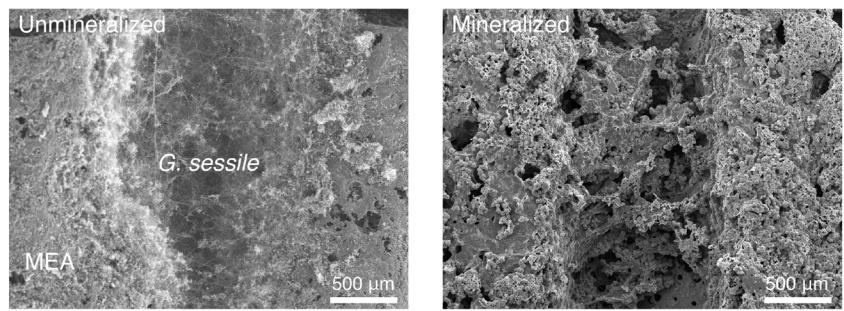

**Fig. S9.**

**SEM images showing the surface morphology of mycelial scaffolds formed by *G. sessile*, before and after biomineralization.** The unmineralized scaffold (left) contains *G. sessile* in MEA-based ink, while the mineralized scaffold (right) shows mineral coverage and roughened morphology.

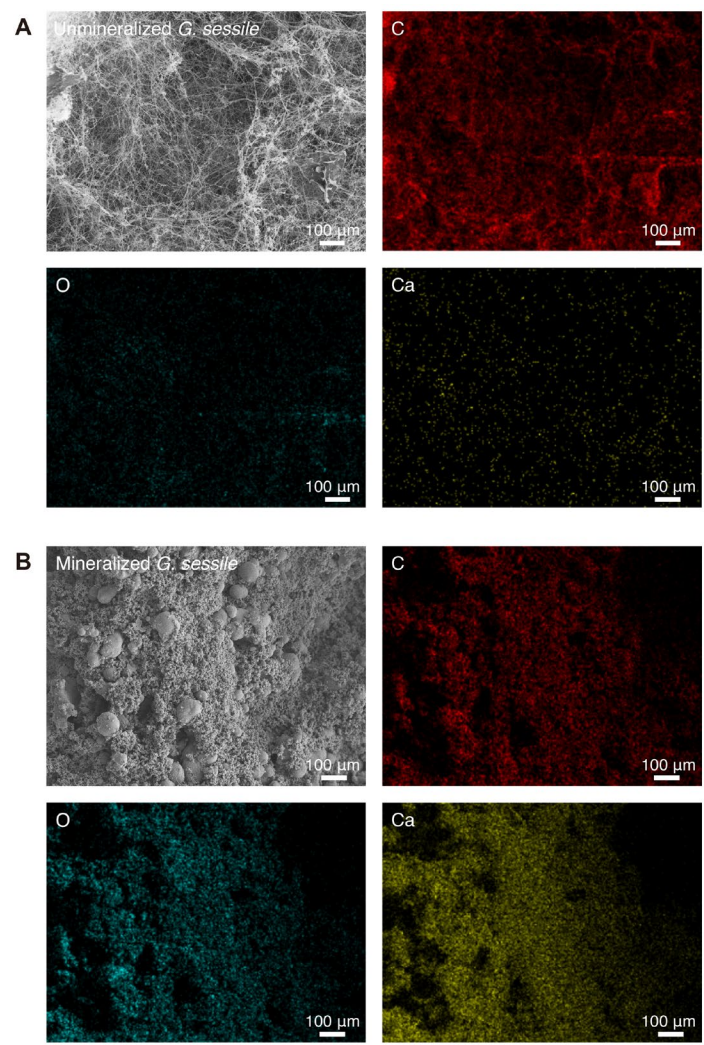

**Fig. S10.**

**Elemental mapping of unmineralized and mineralized mycelial scaffolds formed by *G. sessile*.** (A and B) SEM and EDS elemental mapping of *G. sessile* scaffolds before (A) and after (B) mineralization. Elemental maps show carbon (C, red), oxygen (O, cyan), and calcium (Ca, yellow). The mineralized scaffold exhibits the presence of calcium-based mineral deposits.

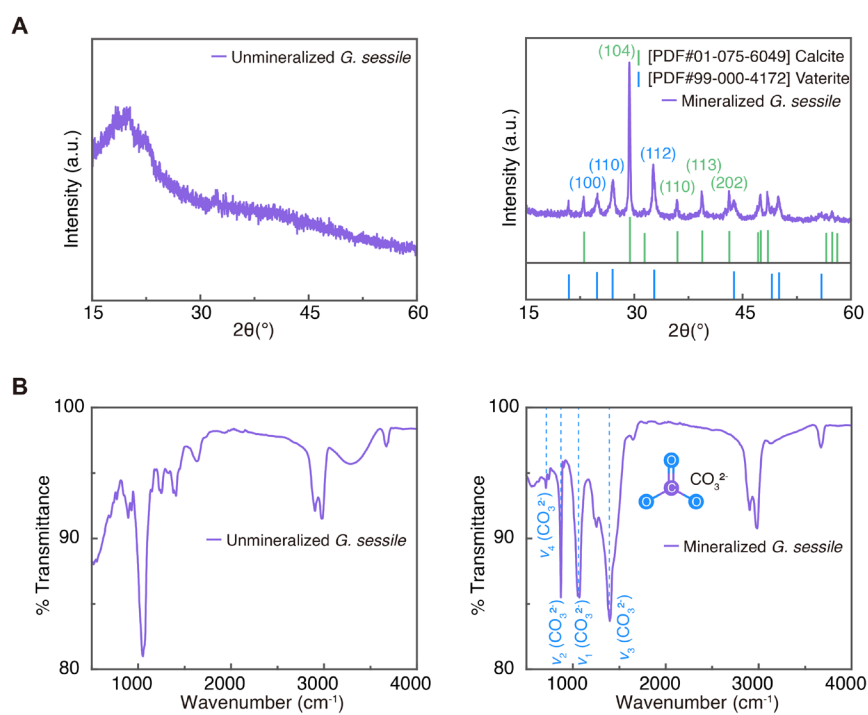

**Fig. S11.**

**PXRD and FTIR characterization of *G. sessile* mycelium scaffolds before and after mineralization.** (A) PXRD patterns of *G. sessile* scaffolds. The unmineralized scaffold (left) shows a broad amorphous peak, while the mineralized scaffold (right) exhibits sharp diffraction peaks corresponding to calcite and vaterite, indicating the formation of crystalline calcium carbonate phases. (B) FTIR spectra of *G. sessile* scaffolds. The unmineralized scaffold (left) exhibits typical absorption bands associated with fungal polysaccharides and proteins. The mineralized scaffold (right) shows four characteristic carbonate absorption bands: $\nu_1$ ($\sim 1070\text{ cm}^{-1}$, symmetric stretching), $\nu_2$ ($\sim 870\text{ cm}^{-1}$, out-of-plane bending), $\nu_3$ ($\sim 1400\text{ cm}^{-1}$, asymmetric stretching), and $\nu_4$ ($\sim 710\text{ cm}^{-1}$, in-plane bending). These bands indicate the presence of carbonate groups ($\text{CO}_3^{2-}$) (2), and when considered together with XRD results, support the deposition of calcium carbonate within the scaffold.

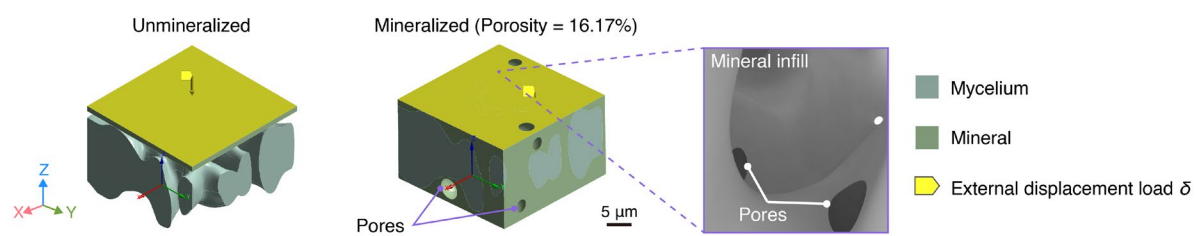

**Fig. S12.**

**Finite element modeling and simulation of unmineralized and mineralized microscale mycelial networks.** Finite element simulations of unmineralized and mineralized mycelial subvolumes under external displacement loading. Mineralization was modeled by partially filling hyphal gaps with calcium carbonate to mimic mineral infiltration, resulting in an average porosity of 16.17%. The porosity value was determined based on microCT image slices.

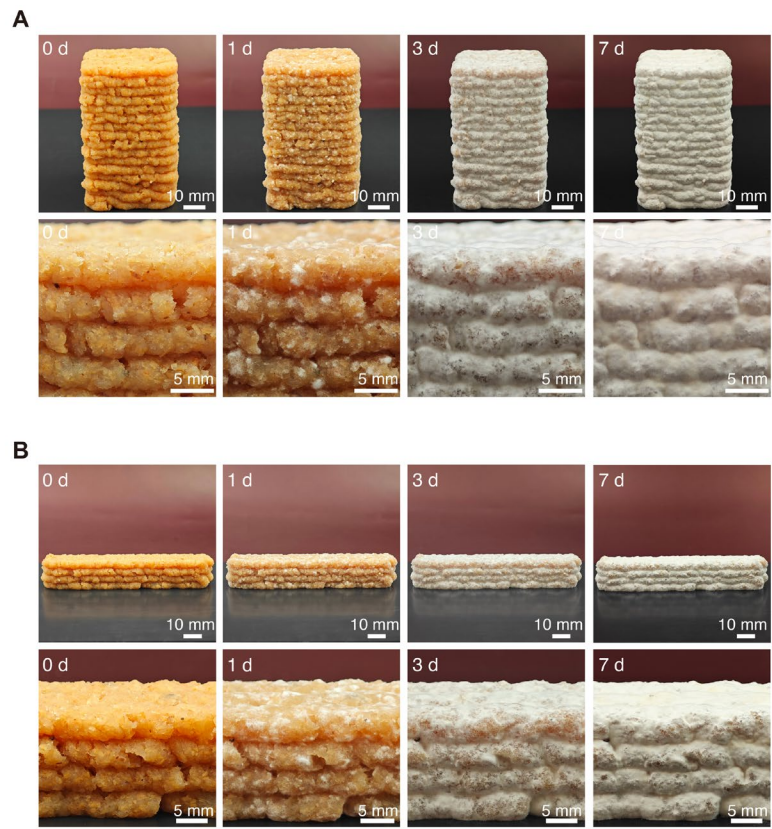

**Fig. S13.**

**Growth of *T. versicolor* in 3D-printed sawdust-based scaffolds for mechanical testing.** (A and B) Overview (top row) and zoom-in views (bottom row) of scaffolds for compression (A) and three-point bending tests (B) over 7 days. Progressive mycelial growth and surface whitening are observed during incubation.

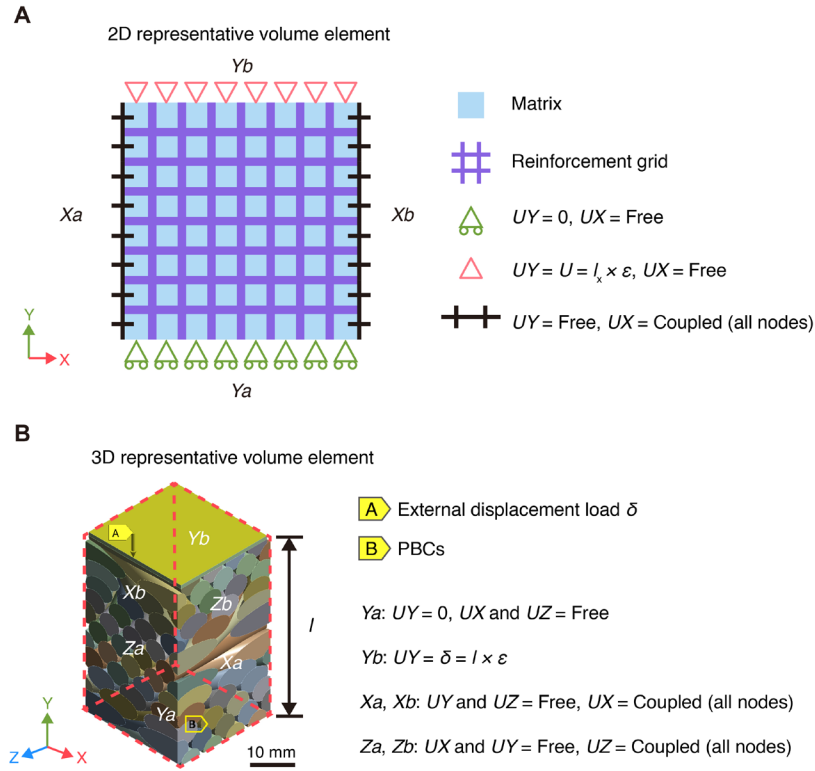

**Fig. S14.**

**PBCs applied to 2D and 3D RVEs of sawdust-based mycelium composites with Bouligand structure under compression. (A) 2D RVE schematic of the mycelium composite. (B) 3D RVE representing the Bouligand structure used in mechanical simulation.**

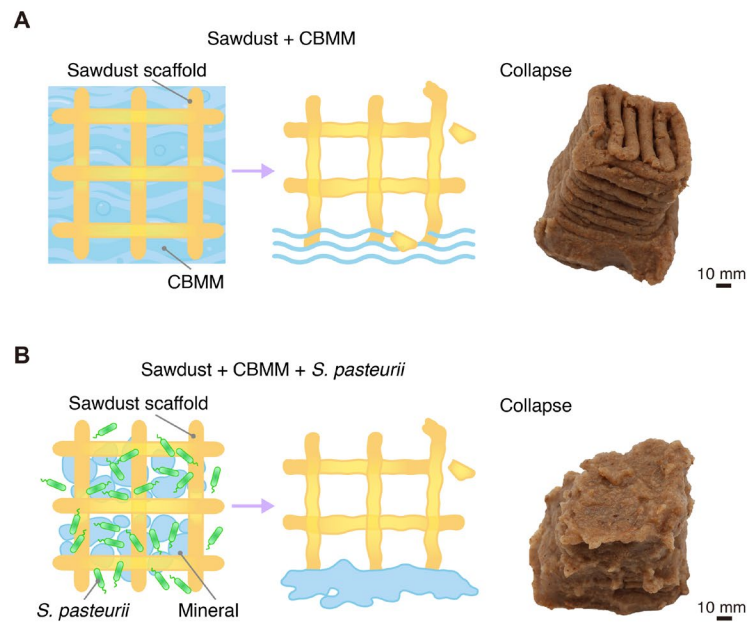

**Fig. S15.**

**Control sawdust-based scaffolds without mycelium show structural collapse after CBMM soaking, with or without *S. pasteurii*.** (A and B) Schematic and photographic illustrations of 3D-printed sawdust scaffolds immersed in CBMM without (A) and with (B) *S. pasteurii*. In both cases, the scaffolds collapse after immersion. Although *S. pasteurii* is present, the absence of mycelium prevents calcium carbonate from depositing onto the scaffold structure, resulting in insufficient mineral reinforcement.

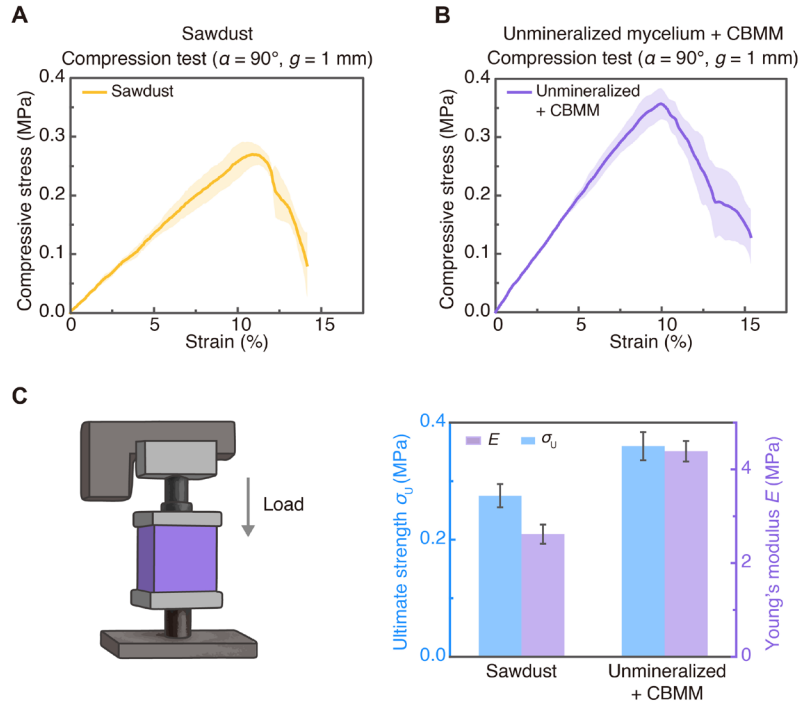

**Fig. S16.**

**Compression test of control scaffolds without mineralization under 90° Bouligand structure ($n = 5$).** (A) Compressive stress–strain curves of 3D-printed sawdust-only scaffold (no mycelium or mineralization), with dark lines as interpolated means, light shading as mean ± SD (upper = mean + SD, lower = mean - SD). (B) Compressive stress–strain curves of unmineralized *T. versicolor* mycelium scaffold after soaking in CBMM, without *S. pasteurii*. Both groups share the same printing parameters: interlayer deflection angle $\alpha = 90^\circ$, line gap $g = 1$ mm. (C) Ultimate compressive strength and Young's modulus of both groups.

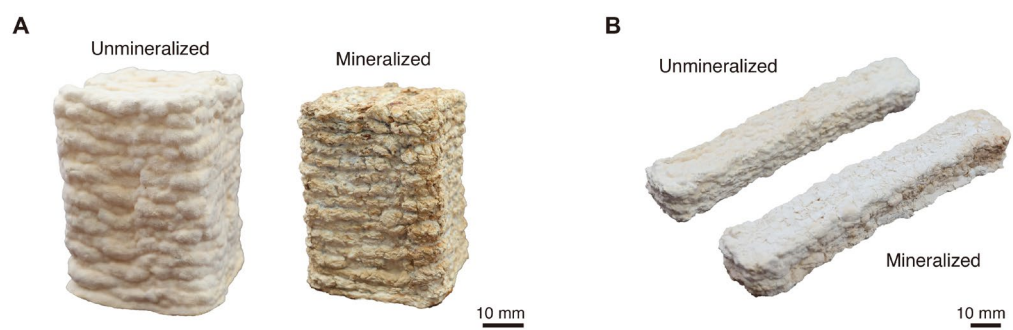

**Fig. S17.**

**Photographs of unmineralized and biomineralized mycelium scaffolds prepared for mechanical testing. (A) Rectangular scaffolds used for compression testing. (B) Beam-shaped mycelium scaffolds used for three-point bending tests.**

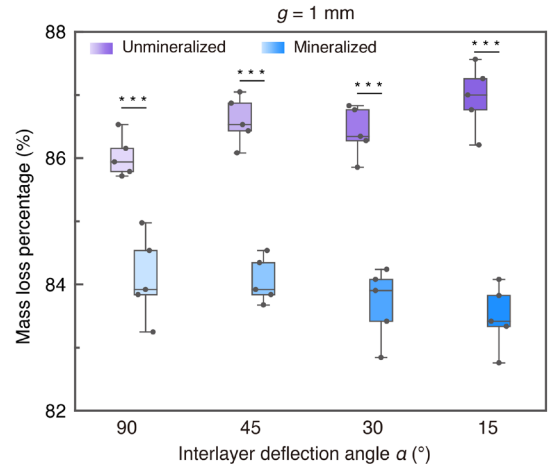

**Fig. S18.**

**Mass loss percentage of *T. versicolor* scaffolds with Bouligand structures of varying interlayer deflection angles after drying ($n = 5$).** Mass loss percentage of unmineralized and mineralized 3D-printed sawdust-based mycelium scaffolds for compression testing, with interlayer deflection angles ($\alpha$) of 90°, 45°, 30°, and 15° (line gap $g = 1$ mm). Mass was recorded before and after drying to calculate the percentage loss (\*$p < 0.05$, \*\*$p < 0.01$, \*\*\*$p < 0.001$, *n.s.*: No significant difference).

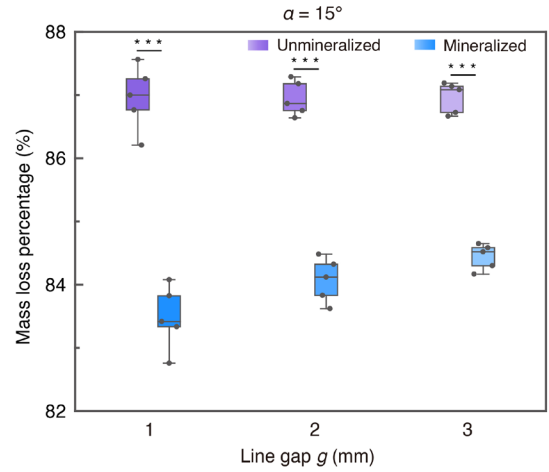

**Fig. S19.**

**Mass loss percentage of *T. versicolor* scaffolds with Bouligand structures at $\alpha = 15^\circ$ and varying line gaps after drying ($n = 5$).** Mass loss percentage of unmineralized and mineralized 3D-printed sawdust-based mycelium scaffolds with line gaps ($g$) of 1 mm, 2 mm, and 3 mm, measured before and after drying (\*$p < 0.05$, \*\*$p < 0.01$, \*\*\*$p < 0.001$, *n.s.*: No significant difference).

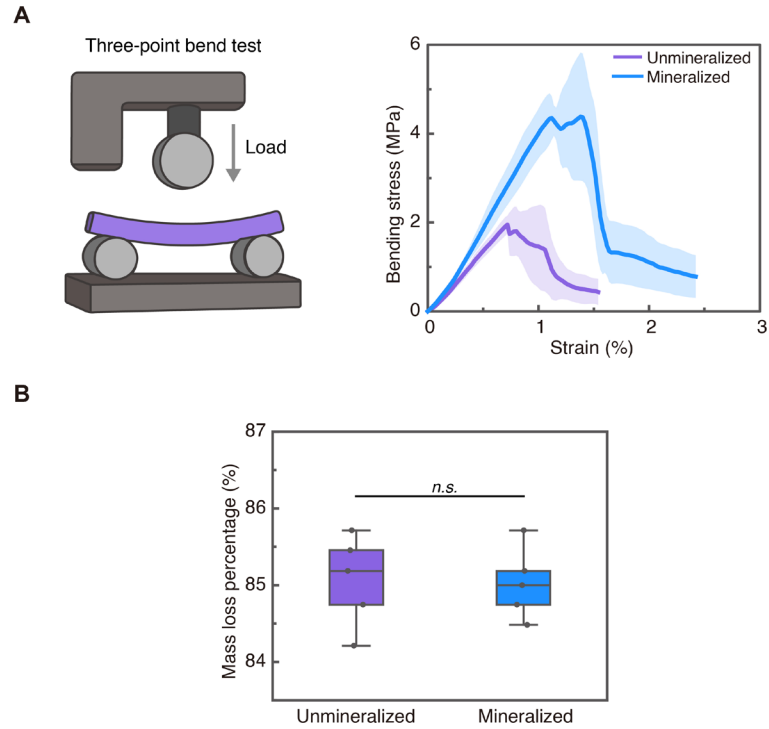

**Fig. S20.**

**Bending performance and mass loss percentage of *T. versicolor* scaffolds under three-point bending tests ($n = 5$).** (A) Bending stress–strain curves of unmineralized (purple) and mineralized (blue) 3D-printed sawdust-based mycelium scaffolds subjected to three-point bending, with dark lines as interpolated means, light shading as mean ± SD (upper = mean + SD, lower = mean - SD). (B) Mass loss percentage of the scaffolds measured before and after drying. No statistically significant difference was observed between the unmineralized and mineralized groups ($*p < 0.05$, $**p < 0.01$, $***p < 0.001$, $n.s.$: No significant difference).

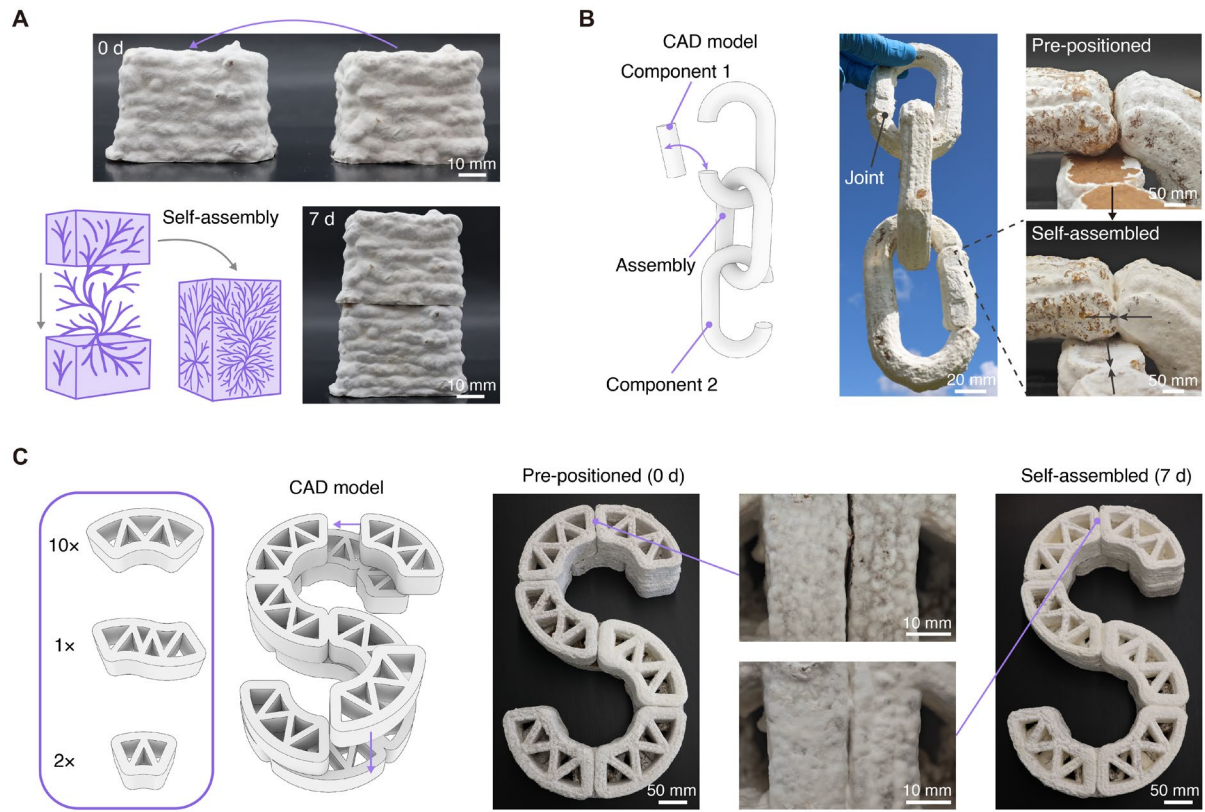

**Fig. S21.**

**Self-assembly of 3D-printed mycelium-based components into integrated structures. (A)** Two mycelium scaffolds of identical size after 7 days of growth brought into contact and subsequently bridged by hyphal extension, forming a continuous structure ($\alpha = 90^\circ$, $g = 1$ mm). **(B)** A chain-like linkage composed of pre-positioned printed modules that were bridged by hyphal growth, transitioning from initial contact to a self-assembled movable joint. **(C)** Modular “S”-shaped structure assembled from discrete mycelium components after 7 days of growth. Hyphal bridging subsequently integrated the modules, resulting in the transition from pre-positioned (0 d) to a connected form (7 d). The pre-positioned (0 d) photograph corresponds to the representative image shown in Fig. 5.

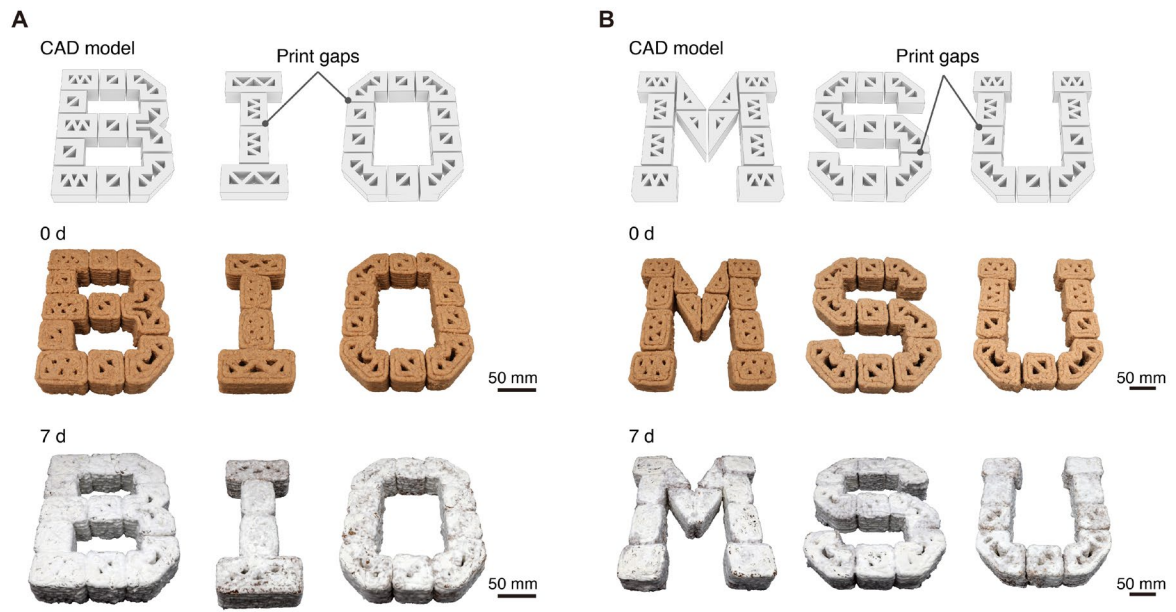

**Fig. S22.**

**Demonstration of self-assembly in 3D-printed mycelium modules.** (A and B) Computer-aided design (CAD) models of the letters "BIO" (A) and "MSU" (B) were printed as discrete modules with print gaps. During 7 days of incubation, the modules were bridged by hyphal growth across the gaps, resulting in the self-assembly of continuous letter structures.

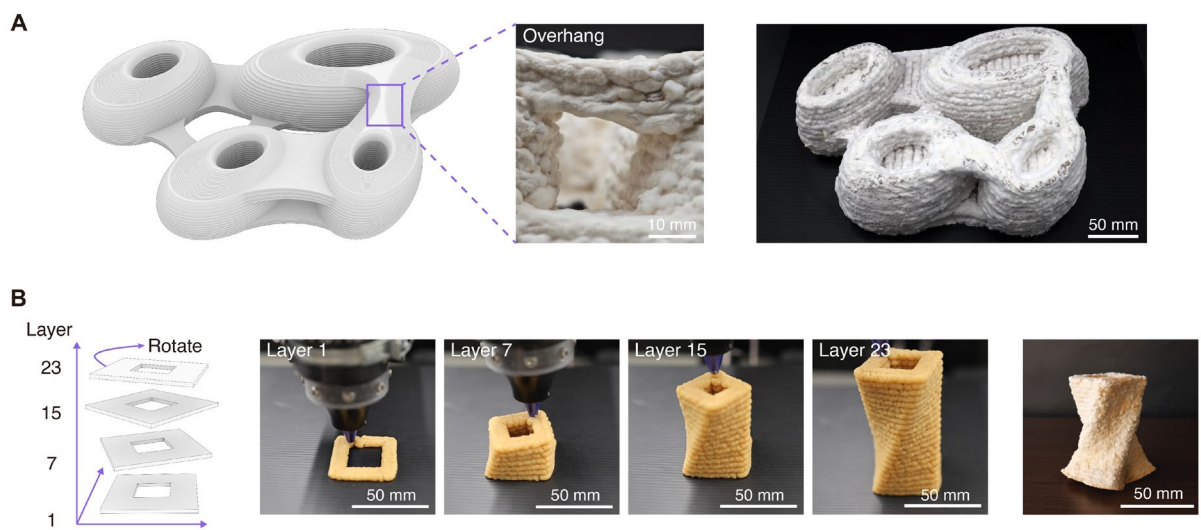

**Fig. S23.**

**Demonstration of 3D printing with sawdust-based mycelium ink.** (A) A recreation of the Galaxy SOHO structure printed with sawdust-based mycelium ink, showing its overhanging details. (B) Printing process of a twisted vase-like structure, showing both schematics and photographs of layers 1, 7, 15, and 23.

### Supplementary Tables

**Table S1.**

Compressive mechanical properties of 3D-printed *T. versicolor* scaffolds with Bouligand structures of varying interlayer angles ($\alpha = 90^\circ, 45^\circ, 30^\circ, 15^\circ$).

| Group | Unmineralized |  |  |  | Mineralized |  |  |  |
| --- | --- | --- | --- | --- | --- | --- | --- | --- |
| $\alpha$ ($^\circ$) | 90 | 45 | 30 | 15 | 90 | 45 | 30 | 15 |
| $g$ (mm) | 1 | 1 | 1 | 1 | 1 | 1 | 1 | 1 |
| $\bar{E}$ (MPa) | 4.38 | 9.37 | 10.23 | 13.16 | 61.23 | 77.20 | 83.38 | 92.18 |
| SD ($E$, MPa) | 0.59 | 1.27 | 1.00 | 1.12 | 7.09 | 2.99 | 4.69 | 3.28 |
| *RSD ($E$, %) | 13.46 | 13.59 | 9.75 | 8.52 | 11.59 | 3.88 | 5.63 | 3.56 |
| $\bar{\sigma}_U$ (MPa) | 0.37 | 0.66 | 0.77 | 0.84 | 1.98 | 3.34 | 3.90 | 4.26 |
| SD ($\sigma_U$, MPa) | 0.02 | 0.03 | 0.05 | 0.03 | 0.06 | 0.18 | 0.18 | 0.18 |
| *RSD ($\sigma_U$, %) | 6.65 | 4.19 | 6.18 | 3.12 | 3.11 | 5.29 | 4.55 | 4.11 |
| $\bar{\rho}$ (g/cm<sup>3</sup>) | 0.29 | 0.32 | 0.33 | 0.33 | 0.42 | 0.46 | 0.47 | 0.47 |
| $E/\rho$ (J/g) | 15.06 | 28.84 | 31.25 | 39.70 | 145.87 | 166.62 | 177.78 | 196.06 |
| $\sigma_U/\rho$ (J/g) | 1.28 | 2.03 | 2.33 | 2.55 | 4.71 | 7.20 | 8.32 | 9.08 |

\* Relative standard deviation (RSD) = Standard deviation (SD) / Mean × 100%

$E$, $\sigma_U$, $\rho$, $E/\rho$, and $\sigma_U/\rho$ are rounded to two decimal places. RSD is reported as percentages and rounded to two decimal places.

**Table S2.**

Compressive mechanical properties of *T. versicolor* scaffolds with Bouligand structures at a fixed interlayer deflection angle ($\alpha = 15^\circ$) and varying line gaps ($g = 2$ mm and 3 mm).

| Group | Unmineralized |  | Mineralized |  |
| --- | --- | --- | --- | --- |
| $\alpha$ (°) | 15 | 15 | 15 | 15 |
| $g$ (mm) | 2 | 3 | 2 | 3 |
| $\bar{E}$ (MPa) | 7.76 | 3.98 | 64.68 | 38.39 |
| SD ($E$, MPa) | 0.31 | 0.35 | 2.74 | 3.30 |
| RSD ($E$, %) | 4.00 | 8.69 | 4.24 | 8.60 |
| $\bar{\sigma}_U$ (MPa) | 0.74 | 0.33 | 3.22 | 1.55 |
| SD ($\sigma_U$, MPa) | 0.03 | 0.02 | 0.17 | 0.12 |
| RSD ($\sigma_U$, %) | 3.60 | 5.61 | 5.14 | 7.77 |
| $\bar{\rho}$ (g/cm<sup>3</sup>) | 0.33 | 0.28 | 0.43 | 0.35 |
| $E/\rho$ (J/g) | 23.84 | 14.48 | 150.54 | 108.26 |
| $\sigma_U/\rho$ (J/g) | 2.28 | 1.19 | 7.50 | 4.37 |

The data for $g = 1$ mm are identical to those presented in Table S1 and are not repeated here for clarity.

**Table S3.**

Compressive mechanical properties of control scaffolds without mineralization ($\alpha = 90^\circ$, $g = 1$ mm).

| Group | Sawdust | Unmineralized mycelium + CBMM |
| --- | --- | --- |
| $\alpha$ ($^\circ$) | 90 | 90 |
| $g$ (mm) | 1 | 1 |
| $\bar{E}$ (MPa) | 2.62 | 4.39 |
| SD ($E$, MPa) | 0.21 | 0.22 |
| RSD ($E$, %) | 7.86 | 5.02 |
| $\bar{\sigma}_U$ (MPa) | 0.28 | 0.36 |
| SD ($\sigma_U$, MPa) | 0.02 | 0.02 |
| RSD ($\sigma_U$, %) | 6.72 | 6.70 |
| $\bar{\rho}$ (g/cm<sup>3</sup>) | 0.22 | 0.30 |
| $E/\rho$ (J/g) | 12.18 | 14.84 |
| $\sigma_U/\rho$ (J/g) | 1.28 | 1.22 |

**Table S4.**

Flexural mechanical properties of *T. versicolor* scaffolds under three-point bending.

| Group | Unmineralized | Mineralized |
| --- | --- | --- |
| $\overline{E_f}$ (MPa) | 282.43 | 373.99 |
| SD ($E_f$, MPa) | 33.07 | 44.67 |
| RSD ($E_f$, %) | 11.71 | 11.94 |
| $\overline{\sigma_U}$ (MPa) | 2.26 | 5.10 |
| SD ($\sigma_U$, MPa) | 0.11 | 0.42 |
| RSD ($\sigma_U$, %) | 4.77 | 8.26 |
| $\overline{\rho}$ (g/cm<sup>3</sup>) | 0.41 | 0.47 |

**Table S5.**

Compressive mechanical properties of self-healing and self-assembly *T. versicolor* scaffolds ($\alpha = 90^\circ$, $g = 1$ mm).

| Group | Self-healing |  | Self-assembly |  |
| --- | --- | --- | --- | --- |
| Treatment | Unmineralized | Mineralized | Unmineralized | Mineralized |
| $\bar{E}$ (MPa) | 4.54 | 56.88 | 4.49 | 55.05 |
| SD ($E$, MPa) | 0.36 | 3.18 | 0.33 | 3.39 |
| RSD ($E$, %) | 8.01 | 5.60 | 7.25 | 6.17 |
| $\bar{\sigma}_U$ (MPa) | 0.38 | 1.79 | 0.33 | 1.79 |
| SD ($\sigma_U$, MPa) | 0.04 | 0.10 | 0.032 | 0.02 |
| RSD ($\sigma_U$, %) | 9.48 | 5.30 | 9.85 | 1.06 |
| $\bar{\rho}$ (g/cm<sup>3</sup>) | 0.31 | 0.41 | 0.32 | 0.41 |
| $E/\rho$ (J/g) | 14.84 | 140.20 | 13.91 | 133.99 |
| $\sigma_U/\rho$ (J/g) | 1.23 | 4.41 | 1.01 | 4.34 |

### **Movies**

#### **Movie S1.**

**3D printing of an Eiffel Tower model using MEA-based mycelium ink.** The video shows the extrusion printing process and shape fidelity of the resulting structure.

#### **Movie S2.**

**Mycelium growth on a 3D-printed "S" structure.** Time-lapse images acquired every 5 min show hyphal colonization and expansion over 7 days.

#### **Movie S3.**

**3D printing of parallel filaments with varying spacing.** The video shows extrusion printing of parallel filaments with controlled spacing for subsequent hyphal bridging experiments.

#### **Movie S4.**

**Compression testing of mycelium scaffolds before and after mineralization.** The video compares the deformation of unmineralized and mineralized scaffolds under compressive loading.

#### **Movie S5.**

**Bending testing of mycelium scaffolds before and after mineralization.** The video compares the mechanical responses of unmineralized and mineralized scaffolds during three-point bending.

#### **Movie S6.**

**Mycelium self-healing.** A single artificial perforation was introduced, and hyphal growth over 4 days demonstrates the repair of the structural discontinuity.

#### **Movie S7.**

**3D printing of a twisted vase using sawdust-based mycelium ink.** The video shows the fabrication of a twisted vase through progressive rotation of successive printed layers.

#### **Movie S8.**

**3D printing of a mycelium dome on a pre-printed support.** The video shows the deposition of mycelium ink onto a pre-printed PLA support and the resulting freestanding hollow dome after support removal.

#### Supplementary Reference:

1. S. Gantenbein, E. Colucci, J. Käch, E. Trachsel, F. B. Coulter, P. A. Rühs, K. Masania, A. R. Studart, Three-dimensional printing of mycelium hydrogels into living complex materials. *Nat. Mater.* **22**, 128–134 (2023).
2. F. Natalio, R. Maria, Microbial biomineralization of alkaline earth metal carbonates on 3D-printed surfaces. *ACS Appl. Mater. Interfaces* **16**, 6327–6336 (2024).
3. E. Viles, E. Heyneman, S. Lin, V. Montague, A. Darabi, L. M. Cox, A. Phillips, R. Gerlach, E. J. Espinosa-Ortiz, C. Heveran, Mycelium as a scaffold for biomineralized engineered living materials. *Cell Rep. Phys. Sci.* **6** (2025).
4. I. M. Van Meerbeek, J. M. Lenhardt, W. Small, T. M. Bryson, E. B. Duoss, T. H. Weisgraber, Compressive properties of silicone Bouligand structures. *MRS Bull.* **48**, 325–331 (2023).
5. M. K. Elhachimi, A. Solhtalab, M. J. Razavi, D. L. Porter, Computational modeling and analysis of fungi-inspired network systems. *Adv. Eng. Mater.*, doi: 10.1002/adem.202402949 (2025).
6. C. M. Heveran, L. Liang, A. Nagarajan, M. Hubler, R. Gill, J. Cameron, S. M. Cook, W. Srubar, Engineered ureolytic microorganisms can tailor the morphology and nanomechanical properties of microbial-precipitated calcium carbonate. *Sci. Rep.* **9** (2019).
7. H.-J. Shin, H.-S. Ro, M. Kawauchi, Y. Honda, Review on mushroom mycelium-based products and their production process: from upstream to downstream. *Bioresour. Bioprocess.* **12**, 3 (2025).
8. M. P. Jones, T. Huynh, S. John, Inherent species characteristic influence and growth performance assessment for mycelium composite applications. *Advanced Materials Letters* **9**, 71–80 (2018).
9. M. P. Jones, A. C. Lawrie, T. T. Huynh, P. D. Morrison, A. Mautner, A. Bismarck, S. John, Agricultural by-product suitability for the production of chitinous composites and nanofibers. *Process Biochem.* **80**, 95–102 (2019).
10. W. C. Oliver, G. M. Pharr, An improved technique for determining hardness and elastic modulus using load and displacement sensing indentation experiments. *J. Mater. Res.* **7**, 1564–1583 (1992).
11. P. A. Amorim, M. A. d'Ávila, R. Anand, P. Moldenaers, P. Van Puyvelde, V. Bloemen, Insights on shear rheology of inks for extrusion-based 3D bioprinting. *Bioprinting* **22**, e00129 (2021).
12. N. Paxton, W. Smolan, T. Böck, F. Melchels, J. Groll, T. Jungst, Proposal to assess printability of bioinks for extrusion-based bioprinting and evaluation of rheological properties governing bioprintability. *Biofabrication* **9**, 044107 (2017).
13. M. R. Islam, G. Tudryn, R. Bucinell, L. Schadler, R. C. Picu, Morphology and mechanics of fungal mycelium. *Sci. Rep.* **7** (2017).
14. M. R. Islam, G. Tudryn, R. Bucinell, L. Schadler, R. C. Picu, Mechanical behavior of mycelium-based particulate composites. *J. Mater. Sci.* **53**, 16371–16382 (2018).
15. M. A. Shakir, B. Azahari, Y. Yusup, M. F. Yhaya, A. Salehabadi, M. I. Ahmad, Preparation and characterization of mycelium as a bio-matrix in fabrication of bio-composite. *Adv. Res. Fluid Mech. Therm. Sci.* **65**, 253–263 (2020).
